# Identification of Reductases Catalyzing Benzyl Alcohol Formation during Salicylic Acid Biosynthesis in Plants

**DOI:** 10.64898/2026.05.13.724993

**Authors:** Lu Xu, Mingsong Wu, Dan Qiu, Josh Li, Chenying Li, Yanan Liu, Xin Li, Yuelin Zhang

## Abstract

Salicylic acid (SA), a central hormone in plant immunity, is biosynthesized via a recently elucidated phenylalanine-derived pathway in most seed plants. This pathway requires benzyl alcohol as a key substrate for the formation of the SA precursor benzyl benzoate. However, how benzyl alcohol is produced in plants was unclear. Here, we identify a two-step conversion of benzoyl-CoA to benzyl alcohol via benzaldehyde in *Nicotiana* (*N.*) *benthamiana*. From a forward genetic screen for SA-deficient mutants, the α and β subunits of heterodimeric benzaldehyde synthase (BalS) involved in the conversion of benzoyl-CoA to benzaldehyde were found to be required for SA biosynthesis in *N. benthamiana*. Further reverse genetic analysis revealed that the NADPH-dependent benzaldehyde reductase (BalR1) acts downstream of BalS to convert benzaldehyde to benzyl alcohol. Interestingly, *OsBalR1*, but not *OsBalSα* or *OsBalSβ*, is required for maintaining high basal SA levels in rice, suggesting the presence of redundant benzoyl-CoA-reducing activities or alternative biosynthesis routes for benzyl alcohol production. Together, this work defines the missing enzymatic steps in phenylalanine-derived SA pathway and provides insights into the evolutionary diversification of SA production strategies in plants.

## Introduction

Plants rely on a multilayered immune system to detect and defend against pathogens. Pattern-triggered immunity (PTI) is activated by plasma membrane-localized pattern recognition receptors (PRRs), whereas effector-triggered immunity (ETI) is triggered by intracellular nucleotide-binding leucine-rich repeat receptors (NLRs) (Jones et al., 2024). Although PTI and ETI are initiated through distinct recognition mechanisms, they converge on overlapping downstream responses, most notably the accumulation of the defense hormone salicylic acid (SA) (Peng et al., 2018).

SA is a key phytohormone that orchestrates both local and systemic defense responses. For decades, SA biosynthesis was thought to occur predominantly through the isochorismate (ICS) pathway, as defined in the model plant *Arabidopsis* (*A.*) *thaliana*. However, recent studies have uncovered a more broadly conserved phenylalanine (Phe)-derived SA biosynthesis pathway, in which SA is synthesized from benzoyl coenzyme A (benzoyl-CoA) through three consecutive enzymatic steps (Liu et al., 2025; Ma et al., 2025; Wang et al., 2025; Zhu et al., 2025). In this pathway, benzoyl-CoA:benzyl alcohol benzoyl transferase (BEBT) conjugates benzoyl-CoA—originating from the peroxisomal β-oxidation pathway—with benzyl alcohol to form benzyl benzoate. The benzyl benzoate oxidase/benzyl benzoate hydroxylase (BBO/BBH) then catalyzes the oxidation of benzyl benzoate to produce benzyl salicylate, which is subsequently hydrolyzed by benzyl salicylate hydrolase/benzoic salicylate esterase (BSH/BSE) to release SA. This pathway represents the predominant route for SA production in seed plants.

Despite these advances, the enzymatic steps responsible for producing benzyl alcohol, an essential co-substrate for BEBT, remain unclear. Huang et al. (2022) characterized a heterodimeric benzaldehyde synthase (BalS), composed of α and β subunits, that catalyzes benzaldehyde formation in *Petunia* (*P.*) *hybrida*, and demonstrated conserved enzymatic activity in *Arabidopsis*, tomato and almond. However, its role in SA biosynthesis was not established. More recently, Ma et al. (2025) showed that mutating the *BalSα* homolog in poplar resulted in reduced SA accumulation. However, the contribution of the BalS β subunit to SA accumulation was not analyzed in the study. More importantly, the downstream step connecting benzaldehyde to benzyl alcohol has not been defined.

Here, we used both forward and reverse genetic approaches to define the missing enzymatic steps in the upstream portion of the Phe-derived SA biosynthesis pathway in *N. benthamiana*. We demonstrate that both α and β subunits of *Nb*BalS are required for SA biosynthesis, functioning as a heterodimeric reductase that converts benzoyl-CoA to benzaldehyde *in N. benthamiana*. Phylogenetic and structural analyses indicate that this heterodimeric complex originated early during land plant evolution, with orthologs from the liverwort *Marchantia* (*M.*) *polymorpha* retaining functional conservation sufficient to restore SA biosynthesis in *N. benthamiana* mutants. We further identify benzaldehyde reductase 1 (BalR1) as an NADPH-dependent enzyme that catalyzes the subsequent conversion of benzaldehyde to benzyl alcohol, which is required for pathogen-induced SA accumulation in seedlings. Finally, analyses in rice reveal that *Os*BalR1, but not *Os*BalSα or *Os*BalSβ, is required to maintain high basal SA levels, highlighting lineage-specific strategies for SA precursor production.

## Results

### Both BalSα and BalSβ are required for SA biosynthesis in *N. benthamiana* by converting benzoyl-CoA into benzaldehyde

To identify enzymes involved in SA biosynthesis, we previously conducted a forward genetic screen for mutants defective in SA accumulation following *Pseudomonas syringae pv. tomato* (*Pst*) DC3000 infection in *N. benthamiana* (Liu et al., 2025). Several mutant lines, including 4-66, 13-270, and 15-328, exhibited markedly reduced SA levels (Fig.1A) (Liu et al., 2025). To test whether their low-SA phenotype reflected a defect in SA biosynthesis pathway rather than in upstream signaling, we transiently expressed the *Arabidopsis At2g32140*, which encodes a protein with only a Toll Interleukin Receptor (TIR)-domain known to activate SA accumulation independent of pathogen infection (Tian et al., 2021), in the *N. benthamiana* wild type (WT) and mutant plants. These mutants all failed to accumulate SA upon *At2g32140* expression, indicating defects in SA biosynthesis or signaling downstream of TIR activity (Supplementary Fig. S1A).

**Figure 1.**
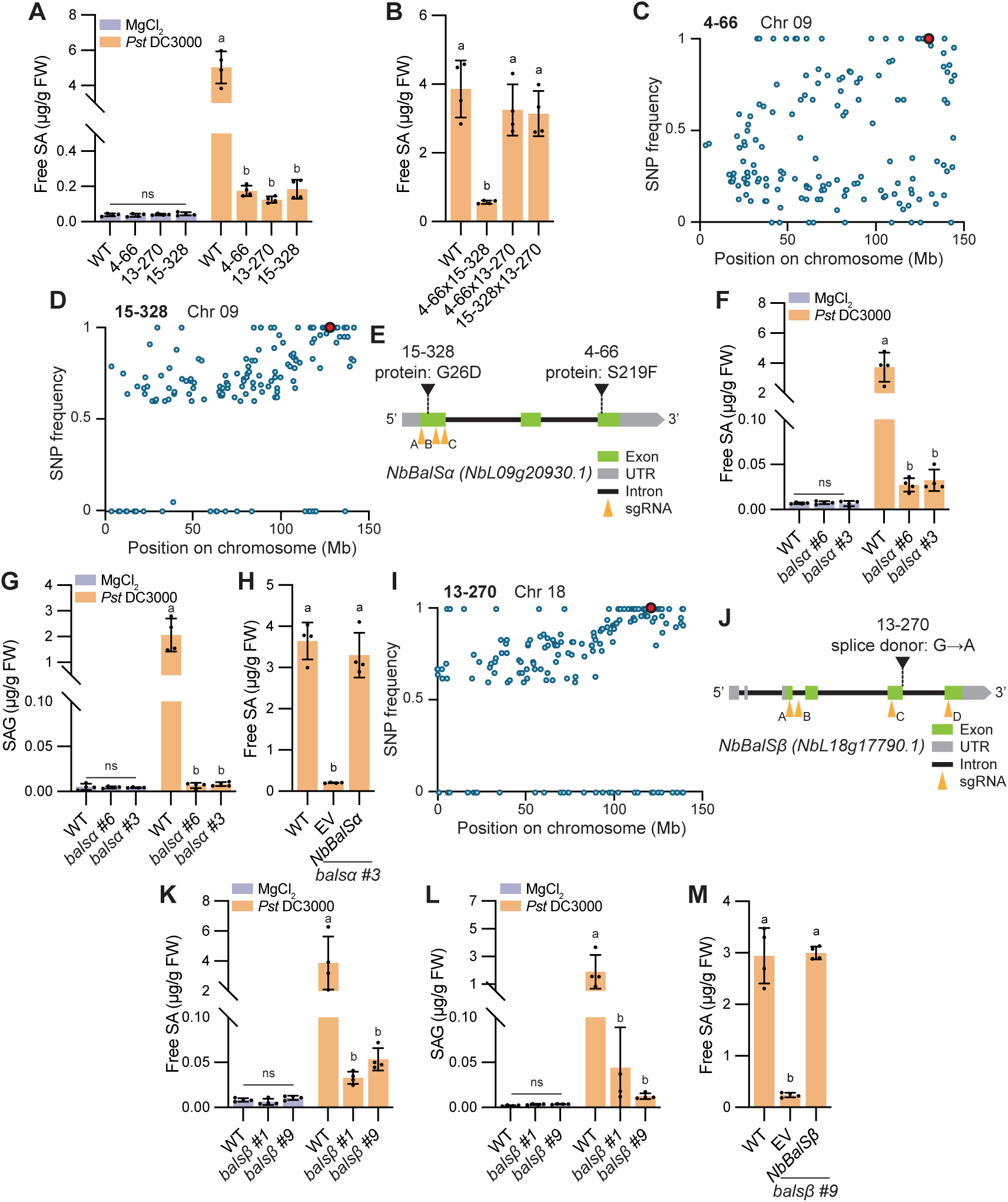
BalSα and BalSβ are required for SA biosynthesis in *N. benthamiana*. **A)** Free SA levels in five-week-old WT and three SA-deficient mutants at 8 hours post inoculation (hpi) with *Pst* DC3000 (OD_600_ = 0.02) or MgCl_2_, as measured using HPLC. **B)** Free SA levels in five-week-old F_1_ hybrids from crosses between the three SA-deficient mutants at 24 hpi with *Pst* DC3000 (OD_600_ = 0.01) measured by a biosensor-based method. **C-D, I)** Single nucleotide polymorphism (SNP) frequency distribution along chromosome 9 (**C-D**) or 18 (**I**) from bulked segregant analysis of SA-deficient individuals pooled from F_2_ populations of mutant lines 4-66 (**C**), 15-328 (**D**) and 13-270 (**I**) with WT, respectively. The red dot marks the candidate *balsα* (**C-D**) or *balsβ* (**I**) mutation with a Percentage of SNP (PSNP) of 100%. Blue dots represent background mutations. Mb, megabase pairs. **E, J)** Structure of the *BalSα* gene (*NbL09g20930.1*) (**E**) or *BalSβ* gene (*NbL18g17790.1*) (**J**), showing the EMS-induced point mutation and CRISPR-Cas9 guide RNA target sites. The mutant 4-66 carries a C-to-T base transition in exon 3, resulting in an S-to-F amino acid substitution. 15-328 has a G-to-A transition in exon 1, causing a G-to-D substitution. 13-270 harbors a G-to-A base transition at the 5’ splice donor site after exon 3, leading to a frameshift. Exons are shown as green rectangles, untranslated regions (UTRs) as grey rectangles, and introns as black lines. Orange triangles denote the protospacer adjacent motif (PAM) sites targeted by sgRNAs. **F-G, K-L)** Free SA (**F, K**) and SAG (**G, L**) levels in five-week-old WT, *balsα* (**F-G**) and *balsβ* (**K-L**) knockout mutants at 8 hpi with *Pst* DC3000 (OD_600_ = 0.02) or MgCl_2_ measured using LC-MS. **H, M)** Free SA levels in five-week-old *balsα* (**H**) and *balsβ* (**M**) CRISPR mutants transiently expressed with empty vector (EV) or *Balsα* and *BalSβ* cDNA, respectively, at 8 hpi with *Pst* DC3000 (OD_600_ = 0.02), as measured using HPLC. Error bars represent mean ± s.d. ns, no significant difference. Letters indicate statistically significant differences (one-way ANOVA with Tukey’s test, n = 4 independent plants).

To identify the causal mutations of SA deficiency in 4-66, 13-270, and 15-328, each mutant was backcrossed with the WT for bulked segregant analysis. The F₁ progeny from two independent crosses for each mutant restored *Pst* DC3000-induced SA accumulation (Supplementary Fig. S1B), and all F₂ populations exhibited an approximately 3:1 segregation ratio of WT to mutant phenotypes (Supplementary Fig. S1C), consistent with the segregation ratio for single recessive mutations. Interestingly, the F₁ progeny from a cross between 4-66 and 15-328 retained low SA levels following *Pst* DC3000 infection, whereas those from crosses between 4-66 and 13-270 or 15-328 and 13-270 did not, suggesting that lines 4-66 and 15-328 carry mutations that are allelic with each other (Fig. 1B). Whole genome sequencing (WGS) of pooled F₂ individuals with the low-SA phenotype (39/261 from 4-66; 50/288 from 15-328) revealed a shared linkage region on chromosome 9 based on single-nucleotide polymorphism (SNP) frequency analysis (Fig. 1C-D). Within this region, 4-66 and 15-328 carried different nonsynonymous substitutions in *NbL09g20930.1*, which encodes a homolog of *P. hybrida* BalSα subunit. 4-66 contains a C-to-T transition causing a Ser219-to-Phe (S219F) substitution, and 15-328 has a G-to-A transition causing a Gly26-to-Asp (G26D) substitution (Fig. 1E; Supplementary Table 1). These findings are consistent with the allelism test results (Fig. 1B) and strongly support that *NbBalSα* is required for SA biosynthesis.

To validate the role of *NbL09g20930.1* in SA biosynthesis, two independent knockout lines were generated using CRISPR-Cas9 (Fig. 1E). Sanger sequencing confirmed that *balsα #3* carries a 7-bp deletion at the sgRNA target site A, a 146-bp deletion between sites B and C, and a G-to-T substitution at site C, whereas *balsα #6* harbors a 246-bp deletion between sites A and B (Supplementary Fig. S1D). Both lines displayed normal morphology (Supplementary Fig. S1E), but abolished *Pst* DC3000-induced accumulation of SA and SA glucoside (SAG), the major conjugated form of SA (Fig. 1F-G). Furthermore, transient expression of *BalSα* cDNA in the CRISPR mutants prior to pathogen infection restored SA accumulation (Fig. 1H), confirming that *BalSα* is required for pathogen-induced SA biosynthesis in *N. benthamiana*.

In parallel, WGS of pooled F₂ individuals of 13-270 with low SA levels (47/236) identified a linkage region on chromosome 18 (Fig. 1I). Within this interval, 13-270 carried a G-to-A substitution at the 5′ splice donor site following the third exon of *NbL18g17790.*1 (cDNA: 522+1G>A), predicted to cause intron retention and a cDNA frameshift (Fig. 1J; Supplementary Table 1). *NbL18g17790.*1 encodes a homolog of *P. hybrida* BalSβ subunit. Two independent *NbBalSβ* knockout lines were generated using CRISPR-Cas9 (Fig. 1J). The *balsβ #1* line contains a 100-bp deletion between target sites A and B, while *balsβ #9* harbours a 1893-bp deletion between target sites B and C (Supplementary Fig. S2A). Both lines abolished *Pst* DC3000-induced SA and SAG accumulations without affecting their morphology (Fig. 1K-L; Supplementary Fig. S2B), and this deficiency was restored by transient expression of *BalSβ* (Fig. 1M). Together, these results demonstrate that both *NbBalSα* and *NbBalSβ* are essential for SA biosynthesis in *N. benthamiana*.

### *Nb*BalS catalyzes benzaldehyde formation

In *P. hybrida*, *Ph*BalSα and *Ph*BalSβ function as a heterodimeric enzyme catalyzing benzaldehyde formation (Huang et al., 2022). *Nb*BalSα and *Nb*BalSβ share high sequence similarity with their *petunia* counterparts (74.5% and 88.3% identity, respectively), prompting us to test whether they have similar enzymatic activity. We expressed and purified recombinant *Nb*BalSα and *Nb*BalSβ from *Escherichia* (*E.*) *coli* (Supplementary Fig. S3A-B). *In vitro* assays showed that neither subunit alone converted benzoyl-CoA to benzaldehyde, whereas benzaldehyde was produced when both proteins were present (Fig. 2A). The Michaelis constant (*K*_m_) for benzoyl-CoA (824.7 µM) was comparable to that reported for *Ph*BalS (Huang et al., 2022) (Supplementary Fig. S3C).

**Figure 2.**
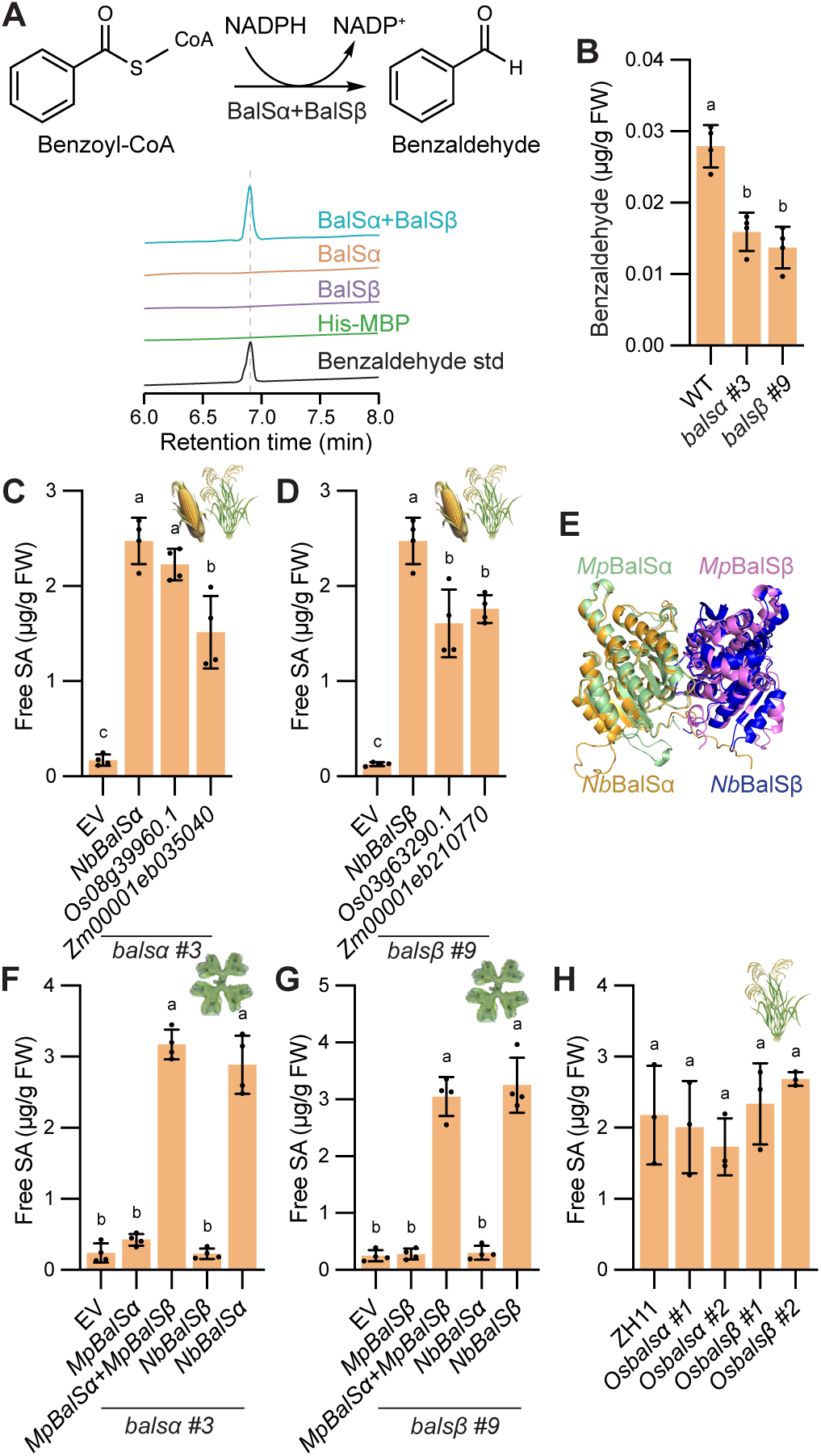
BalS catalyzes benzaldehyde formation and is functionally conserved in plants. **A)** Schematic representation and *in vitro* validation of benzoyl-CoA reduction to benzaldehyde catalyzed by BalSα and BalSβ. Reactions containing both subunits, each subunit alone, or the His-MBP tag control were incubated with benzyl-CoA and NADPH at 28 °C for 30 min. Benzaldehyde formation was verified by HPLC-DAD against an authentic standard. **B)** Levels of benzaldehyde in *NbBEBT*-silenced WT, *balsα* and *balsβ* mutants at 8 hpi with *Pst* DC3000 (OD_600_ = 0.02), as measured using GC-QqQ-MS/MS. *N. benthamiana* leaves were infiltrated with the hpRNA silencing vector targeting *NbBEBT* (OD_600_ = 0.4) three days before *Pst* DC3000 inoculation. **C-D)** Free SA levels in five-week-old *Nbbalsα* (**C**) and *Nbbalsβ* (**D**) mutants transiently overexpressed with their rice or maize orthologs at 8 hpi with *Pst* DC3000 (OD_600_ = 0.02) measured by a biosensor-based method. *N. benthamiana* leaves were infiltrated with *Agrobacterium* carrying the overexpression constructs (OD_600_ = 0.4) three days prior to *Pst* DC3000 inoculation. **E)** Superimposition of the *Mp*BalSα/*Mp*BalSβ (green/purple) and *Nb*BalSα/*Nb*BalSβ (yellow/blue) complexes reveals high similarity in structure, with an overall RMSD of 0.706 Å. Structures were modeled using AlphaFold2 and analyzed using PyMOL. **F-G)** Free SA levels in five-week-old *Nbbalsα* (**F**) and *Nbbalsβ* (**G**) mutants transiently expressing *M. polymorpha* homolog with or without its *M. polymorpha* counterparts at 8 hpi with *Pst* DC3000 (OD_600_ = 0.02) measured by a biosensor-based method. *N. benthamiana* leaves were infiltrated with *Agrobacterium* carrying the overexpression constructs (OD_600_ = 0.4) three days before *Pst* DC3000 inoculation. **H)** Free SA levels in WT, *Osbalsα* and *Osbalsβ* mutants, as measured by LC-MS. Error bars represent mean ± s.d. Letters indicate statistically significant differences (one-way ANOVA with Tukey’s test, n = 4 independent plants).

To assess the *Nb*BalSα and *Nb*BalSβ activity *in planta*, we quantified benzaldehyde levels in mutant and WT plants using gas chromatography-triple quadrupole mass spectrometry (GC-QqQ-MS/MS). Since endogenous benzaldehyde is typically low in *N. benthamiana* leaves (Huang et al., 2022), plants were pretreated with *Pst* DC3000 to stimulate pathway flux. However, benzaldehyde remained barely detectable, likely due to its rapid conversion into downstream metabolites such as benzoic acid (BA) (Long et al., 2009; Saini et al., 2017) and benzyl alcohol (Huang et al., 2022). To suppress the metabolic fluxes and allow benzaldehyde accumulation, we silenced *NbBEBT*, an enzyme catalyzing the third last step toward SA biosynthesis (Liu et al., 2025), using *Agrobacterium*-mediated transient expression of hairpin RNAs targeting *NbBEBT*. Under this condition, benzaldehyde levels were shown to be significantly higher in WT than in *balsα* or *balsβ* mutants (Fig. 2B), supporting an *in vivo* role for *Nb*BalSα and *Nb*BalSβ in benzaldehyde formation. Green fluorescent protein (GFP)-based subcellular localization showed that both subunits localize to the peroxisome (Supplementary Fig. S4A-B), while bimolecular fluorescence complementation (BiFC) assay further confirmed their interaction within this compartment (Supplementary Fig. S4C), in agreement with the peroxisomal origin of the upstream intermediates for Phe-derived SA biosynthesis.

### Cross-species functional conservation of the BalSα-BalSβ complex

Phylogenetic analysis suggests that BalSα and BalSβ emerged in early land plants, with BalSα undergoing lineage-specific expansion and BalSβ remaining predominantly single-copy (Supplementary Fig. S5-7, Supplementary Table 2). To assess whether BalS function is conserved across plant lineages, we performed cross-species complementation assays. Expression of *BalSα* and *BalSβ* orthologs from rice (*Oryza sativa*; *Os08g39960.1* and *Os03g63290.1*) and maize (*Zea mays*; *Zm00001d031453* and *Zm00001d012849*) in *Nbbalsα* and *Nbbalsβ* mutants restored pathogen-induced SA accumulation, either fully or partially (Fig. 2C-D), indicating that monocot enzymes retain their conserved biochemical function.

To examine conservation in early land plants, we analyzed homologs from the liverwort *M. polymorpha*, the earliest lineage containing both BalSα and BalSβ (Supplementary Fig. S5-7). Structural superimposition of *Mp*BalS and *Nb*BalS subunits revealed strong similarity (RMSD = 0.706 Å; Fig. 2E). Expression of *MpBalSα* (*Mp8g05800.1*) or *MpBalSβ* (*Mp5g09150.1*) alone failed to rescue SA accumulation in the corresponding *N. benthamiana* mutants, whereas co-expression of both subunits restored SA production (Fig. 2F-G), suggesting that heterodimer formation is essential and likely established early during land plant evolution.

Despite the ability of rice homologs to complement the *N. benthamiana* mutants (Fig. 2C-D), CRISPR mutants of *OsBalSα* (*Os08g39960.1*) and *OsBalSβ* (*Os03g63290.1*) accumulated SA to WT-like levels (Fig. 2H; Supplementary Fig. S8A), suggesting that rice contains functional BalSα and BalSβ homologs but can generate benzaldehyde through a BalS-independent route. To further assess this evolutionary divergence, we examined *A. thaliana*, which contains six *BalSα* homologs but a single *BalSβ* (*AT3G01980.1*) (Supplementary Fig. S5-7). Consistent with the predominant role of the ICS pathway in this species (Dempsey et al., 2011; Rekhter et al., 2019; Torrens-Spence et al., 2019), the *Atbalsβ* T-DNA mutant showed no obvious reduction in pathogen-induced SA accumulation compared to Col-0 (Supplementary Fig. S8B-D).

To examine the structural basis of *Nb*BalSα-*Nb*BalSβ heterodimerization, we analyzed the predicted inter-subunit interface. Site directed mutagenesis of key residues identified by RING showed that substitution of BalSα Tyr257 (BalSα^Y257A^) weakened interaction with BalSβ, resulting in partial loss of enzymatic activity and *in planta* function (Supplementary Fig. S9A, D-F), whereas the BalSα^E264A/Y265A^ double mutant abolished heterodimer formation, enzymatic activity, and SA restoration (Supplementary Fig. S9B-F). Alignment of these interface regions across species revealed partial conservation, strongest in dicots and more divergent in monocots and *Marchantia* (Supplementary Fig. S9G-H), consistent with the complementation patterns observed earlier (Fig. 2C-D, F-G).

At the structural level, *Nb*BalSα and *Nb*BalSβ exhibit strong symmetry (RMSD = 0.737 Å) (Supplementary Fig. 9I) and highly similar secondary structure organization (Supplementary Fig. S9J). Structural modeling further indicates that overall subunit architecture is conserved from bryophytes to angiosperms, with RMSD values generally below 1 Å relative to the *N. benthamiana* proteins (Supplementary Fig. S5).

### *Nb*BalR1 catalyzes benzaldehyde reduction and contributes to SA biosynthesis during early plant development

To this point, the final unresolved step in SA biosynthesis is the reduction of benzaldehyde to benzyl alcohol. Alcohol dehydrogenases (ADHs), a large enzyme family capable of catalyzing the interconversion of aldehydes and alcohols (Strommer, 2011), represent strong candidates for this reaction. To identify the responsible enzyme, we mined our previous RNA-seq dataset from *Pst* DC3000-infected *N. benthamiana* plants (Liu et al., 2025). Among all annotated ADHs, *NbL12g09650.1* stood out as the most strongly induced gene upon pathogen challenge (Fig. 3A). Notably, analysis of an independent RNA-seq dataset examining gene induction by the bacterial effector XopQ (Qi et al., 2018), a type III effector recognized by the TNL receptor Roq1 to trigger immune responses, identified *NbL12g09650.1* as the only ADH significantly upregulated (Supplementary Table 3), highlighting it as the top candidate.

**Figure 3.**
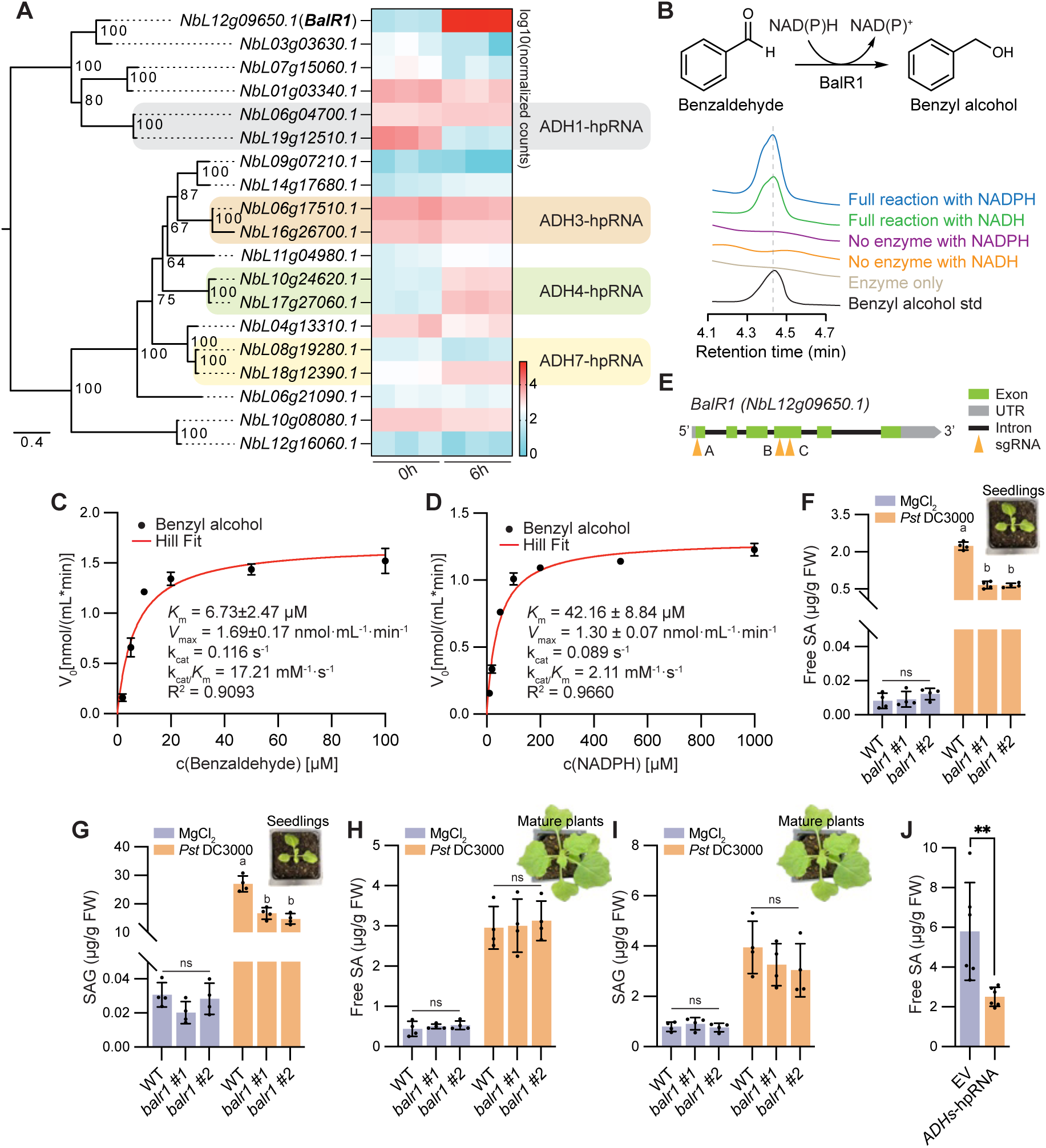
BalR1 catalyzes benzaldehyde reduction and is required for SA accumulation in *N. benthamiana* seedlings. **A)** Phylogenetic tree of alcohol dehydrogenases (ADHs) and heatmap showing *ADH* gene expression at 6 hpi with *Pst* DC3000 (OD_600_ = 0.02) in four-week-old WT *N. benthamiana* plants based on RNA-Seq data (Liu et al., 2025). *ADH* genes targeted by silencing constructs are highlighted. Expression values represent log_10_-transformed normalized read counts derived from DESeq2. **B)** Schematic representation and *in vitro* validation of benzaldehyde reduction to benzyl alcohol catalyzed by BalR1. Reactions with or without BalR1 were incubated with benzaldehyde and NADPH or NADH at 28 °C for 30 min. Benzyl alcohol formation was verified by HPLC-DAD against an authentic standard. RT, retention time. **C-D)** Kinetic curves of BalR1 reaction with varying concentrations of benzaldehyde (**C**) or NADPH (**D**). Data from three replicates were fitted using non-linear Hill regression in Prism 10 (v10.2.0). **E)** Structure of *BalR1* (*NbL12g09650.1*), showing CRISPR-Cas9 guide RNA target sites. Exons are shown as green rectangles, UTRs as grey rectangles, and introns as black lines. Orange triangles denote PAM sites targeted by sgRNAs. **F-I)** Free SA (**F, H**) and SAG (**G, I**) levels in young seedlings (**F-G**) and mature plants (**H-I**) of WT and *balr1* knockout mutants at 8 hpi with *Pst* DC3000 (OD_600_ = 0.02), as measured by LC-MS. **J)** SA levels in *ADH*-silenced plants relative to the empty vector (EV) control at 8 hpi with *Pst* DC3000 (OD_600_ = 0.02) as measured by HPLC. Five-week-old *N. benthamiana* WT leaves were infiltrated with EV or the indicated hpRNA silencing constructs targeting *ADH1/3/4/7* (each at OD_600_ = 0.4) three days prior to *Pst* DC3000 inoculation. hpRNA constructs are color-labeled in (**A**). Error bars represent mean ± s.d. ns, no significant difference. Letters indicate statistically significant differences (one-way ANOVA with Tukey’s test, n = 4 independent plants). Asterisks indicate statistically significant differences (two-tailed Student’s *t*-tests, \**P* < 0.05, n = 4 or 6 independent plants).

To test its enzymatic activity, we expressed and purified the recombinant protein from *E. coli* (Supplementary Fig. S10A). *In vitro* enzymatic assays revealed that it catalyzes the reduction of benzaldehyde to benzyl alcohol using either NADPH or NADH as cofactors (Fig. 3B). Therefore, we designated *NbL12g09650.1* as *benzaldehyde reductase 1* (*BalR1*). Further kinetic analysis revealed a strong preference for NADPH. With NADPH, BalR1 reached a clear plateau, yielding a *K*_m_ of 6.73 µM for benzaldehyde (Fig. 3C) and 42.16 µM for NADPH (Fig. 3D). In contrast, NADH supported detectable activity only at concentrations ≥ 200 µM, with a much weaker affinity (*K*_m_ = 4.94 mM) (Supplementary Fig. S10B). At equivalent cofactor concentrations, initial rates supported by NADH were only 3.54 %, 14.62 %, and 29.71 % of those measured with NADPH at 200, 500, and 1000 µM, respectively. These data indicate that NADPH is likely a preferred cofactor for *Nb*BalR1.

We recently reported that the Phe-derived SA biosynthesis pathway is regulated primarily through the EDS1 (ENHANCED DISEASE SUSCEPTIBILITY 1)-SAG101 (SENESCENCE ASSOCIATED GENE 101)-NRG1 (N REQUIREMENT GENE 1) signaling branch, rather than the EDS1-PAD4 (PHYTOALEXIN DEFICIENT 4)-ADR1 (ACTIVATED DISEASE RESISTANCE 1) module as in *Arabidopsis*, with *BEBT*, *BBO1/2*, and *BSH1/2* losing pathogen-induced expression in *eds1a, sag101b*, and *nrg1* single mutants, but not in *pad4* or *adr1* mutants (Xu et al., 2026). To determine whether *BalR1* follows a similar regulatory pattern, we examined its pathogen-induced expression across the same set of mutants. Notably, *BalR1* exhibited an expression pattern similar to that of established pathway genes, whereas BalSα and BalSβ did not (Supplementary Fig. S10C-E).

To assess the role of *Nb*BalR1 in SA biosynthesis, we generated *Nbbalr1* knockout mutants using CRISPR-Cas9 (Fig. 3E; Supplementary Fig. S10F). Analysis of these mutant lines showed that they exhibited reduced SA and SAG accumulation at the seedling stage (Fig. 3F-G). Notably, this defect was absent in mature plants, which accumulated pathogen-induced SA levels comparable to WT (Fig. 3H-I). This stage-dependent phenotype suggests functional redundancy with other ADH isoforms at later stages. Indeed, silencing additional *ADHs* upregulated by *Pst* DC3000 resulted in reduced pathogen-induced SA accumulation in mature plants (Fig. 3J; Supplementary Fig. S10G). Together, these results indicate that *Nb*BalR1 catalyzes the conversion of benzaldehyde to benzyl alcohol and contributes to SA biosynthesis in *N. benthamiana*, particularly during early development.

We next examined the subcellular localization of *Nb*BalR1. GFP-tagged *NbBalR1* colocalized with the peroxisomal marker mCherry-PTS1 in *N. benthamiana* leaves, indicating its peroxisomal localization (Fig. 4). This localization is consistent with the peroxisomal origin of upstream intermediates and supports its function alongside BalS in sequential pathway steps within this compartment.

**Figure 4.**
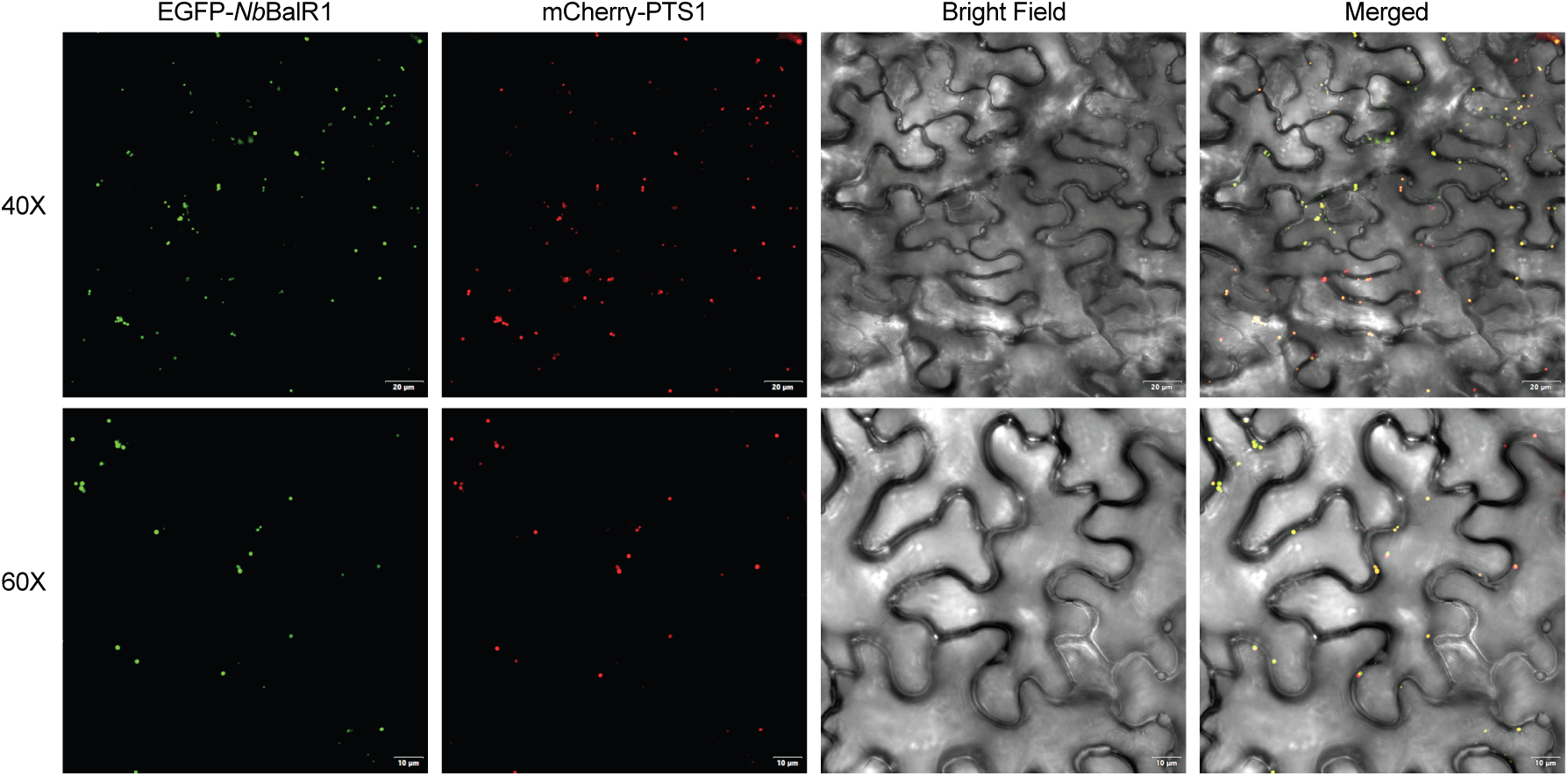
Subcellular localization of EGFP-BalR1 to the peroxisome. EGFP-BalR1 was transiently co-expressed with the peroxisome marker mCherry-PTS1 in five-week-old *N. benthamiana* leaves and visualized by confocal microscopy. EGFP fluorescence (BalR1 fusion protein), mCherry fluorescence (peroxisome), and merged images are shown. Images were acquired at 40× and 60× magnification.

### BalR1 is essential for SA biosynthesis in rice

To assess the conservation of BalR1 function, we analyzed its rice homolog *Os*BalR1 (*Os04g15920.1*). Co-expression analysis using the ATTED-II database (https://atted.jp) showed that *OsBalR1* is co-expressed with the other known SA biosynthetic genes, including *OsCNL*, *OsKAT*, *OsBBO*, *OsBEBT* and *OsPAL7* (Supplementary Table 4). We further generated CRISPR mutants targeting *OsBalR1* (Supplementary Fig. S11A). Notably, *Osbalr1* mutants showed reduction in both SA and SAG levels (Fig. 5A-B), demonstrating that BalR1 is required for SA production in rice. Consistent with this, recombinant *Os*BalR1 purified from *E. coli* catalyzed the reduction of benzaldehyde to benzyl alcohol *in vitro* using NADPH or NADH as cofactors (Fig. 5C; Supplementary Fig. S11B). Similar to *Nb*BalR1, *Os*BalR1 showed a stronger preference for NADPH, yielding a *K*_m_ of 274.58 µM for NADPH and 1.57 mM for benzaldehyde (Fig. 5D-E). In contrast, NADH-supported activity did not reach a plateau at feasible NADH concentrations, indicating much lower affinity and precluding reliable *K*_m_ calculation.

**Figure 5.**
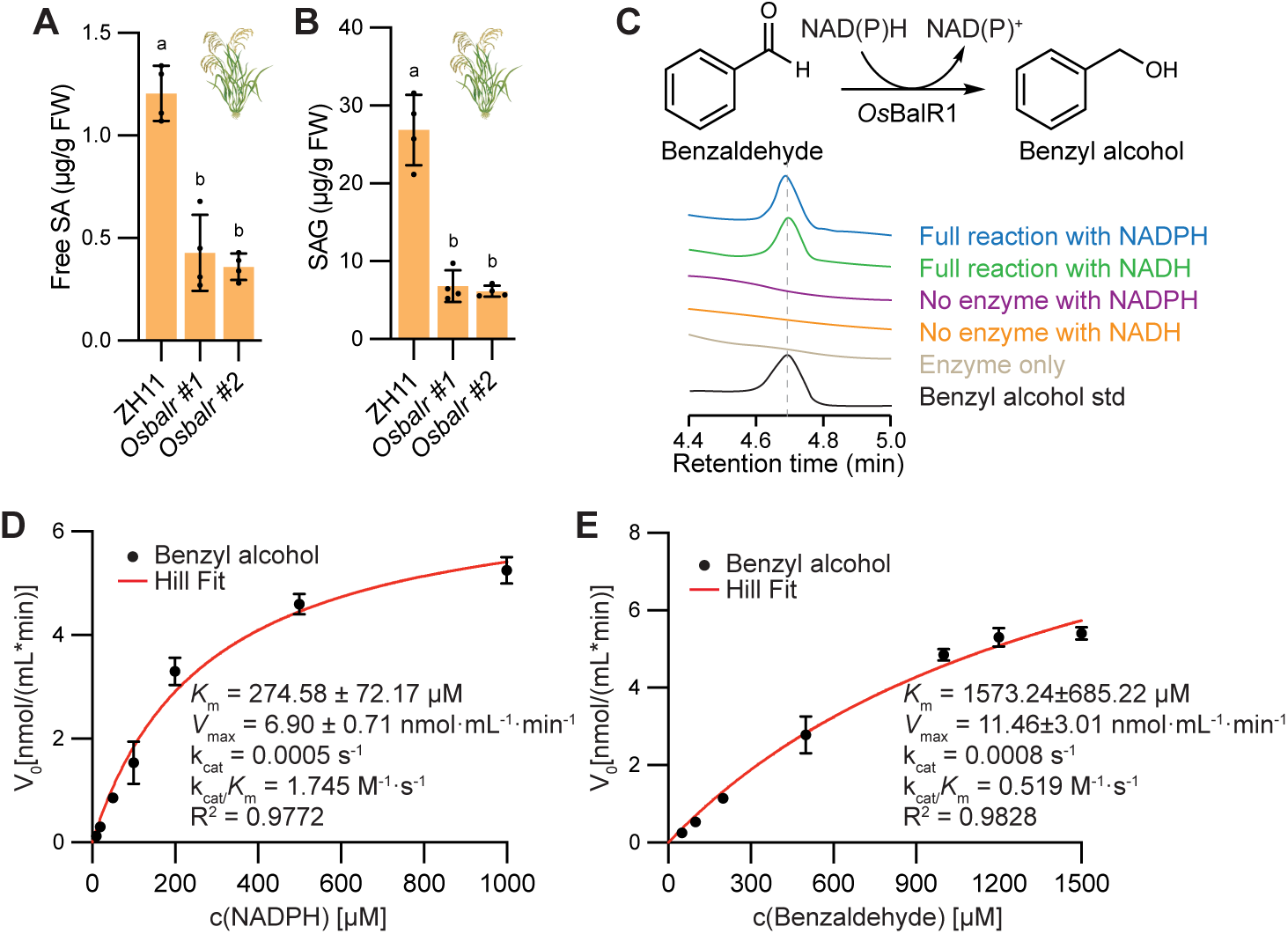
*Os*BalR1 is required for SA accumulation in rice. **A-B)** Free SA (**A**) and SAG (**B**) levels in WT and *Osbalr1* mutants, as measured by LC-MS. Error bars represent mean ± s.d. Letters indicate statistically significant differences (one-way ANOVA with Tukey’s test, n = 4 independent plants). **C)** *In vitro* reduction of benzaldehyde to benzyl alcohol by purified *Os*BalR1. Reactions with or without *Os*BalR1 were incubated with benzaldehyde and NADPH or NADH at 28 °C for 30 min. Benzyl alcohol formation was verified by HPLC-DAD against an authentic standard. RT, retention time. **D-E)** Kinetic curves of *Os*BalR1 reaction with varying concentrations of NADPH (**D**) or benzaldehyde (**E**). Data from three replicates were fitted using non-linear Hill regression in Prism 10 (v10.2.0).

## Discussion

SA is a central phytohormone that orchestrates plant immunity. Recent studies have established the main steps of the Phe-derived SA biosynthesis pathway, in which SA is synthesized from benzoyl-CoA through the sequential actions of BEBT, BBO/BBH, and BSH/BSE (Liu et al., 2025; Ma et al., 2025; Wang et al., 2025; Zhu et al., 2025). Although the origin of benzoyl-CoA has been linked to the β-oxidation pathway, the source of benzyl alcohol—the required co-substrate for BEBT—remained unresolved. Here, we identify BalSα-BalSβ and BalR1 as enzymes that sequentially convert benzoyl-CoA to benzyl alcohol, thereby completing the missing upstream branch of this pathway in *N. benthamiana* (Fig. 6).

**Figure 6.**
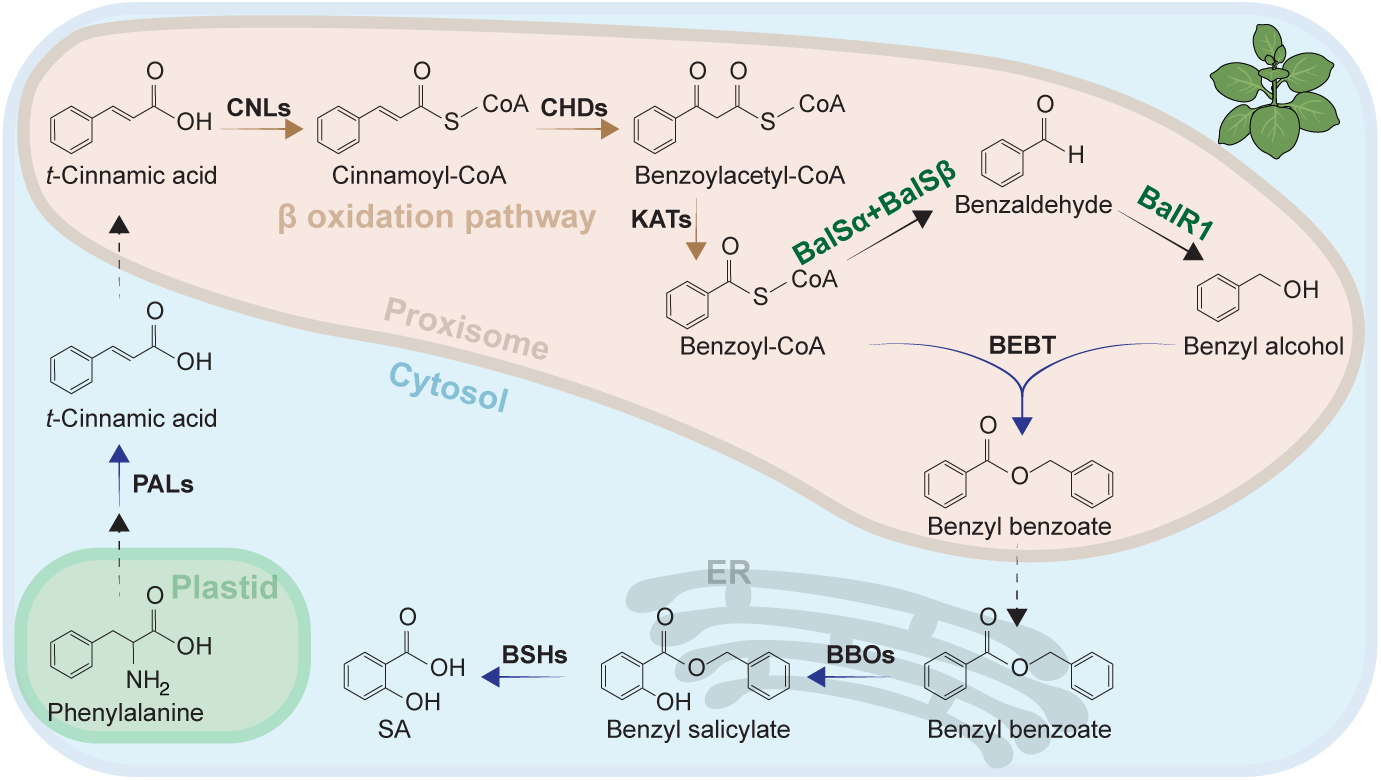
Model of the SA biosynthesis pathway in *N. benthamiana*. The BalSα-BalSβ complex and BalR1 catalyze two steps within the peroxisome, utilizing benzoyl-CoA derived from β-oxidation and producing benzyl alcohol for downstream SA biosynthesis. Both enzymes are shown in dark green font.

BalSα and BalSβ belong to the short-chain dehydrogenase/reductase (SDR) superfamily, a group characterized by low sequence identity (15-30%) and high functional plasticity, which often limits functional prediction from sequences alone (Kallberg et al., 2002; Oppermann et al., 2003). Consistent with this, BalSα and BalSβ exhibit limited sequence conservation yet form an obligate heterodimer essential for catalytic activity (Fig. 2A; Supplementary Fig. S9D-F). Structural analyses reveal that the overall architecture of the BalSα-BalSβ complex is highly conserved across species (Supplementary Fig. S5), despite variations in individual interface residues, suggesting that preservation of structural compatibility underpins enzymatic function. In line with this, orthologs of BalSα and BalSβ from different species can assemble into functional heterodimers (Fig. 2C-D, F-G). However, the partial complementation of SA deficiency observed in the *Nbbalsα* and *Nbbalsβ* mutants observed for some combinations indicates that divergence in primary sequence may modulate the efficiency of heterodimer assembly and activity. The requirement for cognate subunit pairing in *Marchantia* (Fig. 2F-G) further supports co-evolution between the two subunits.

Our study identified BalR1 as a key enzyme in SA biosynthesis. In *N. benthamiana*, loss of *NbBalR1* reduces SA accumulation during early development (Fig. 3F-G), and disruption of *OsBalR1* similarly decreases SA and SAG levels in rice (Fig. 5A-B). Together with *in vitro* enzymatic activity data (Fig. 3B, 5C), these mutant phenotypes support a critical role for BalR1 in the biosynthesis of the SA precursor benzyl alcohol. The reduced SA accumulation observed in *Nbbalr1* mutants at the seedling stage, but not in mature plants, suggests compensation by other ADH isoforms at later stages (Fig. 3F-I), likely enabling flexible control of SA biosynthesis during development. Consistent with this prediction, silencing additional *ADHs* reduced SA levels in mature plants (Fig. 3J).

Consistent with previous reports in *N. benthamiana* and poplar (Huang et al., 2022; Ma et al., 2025), benzaldehyde does not accumulate in WT *N. benthamiana* plants or in mutants lacking the benzaldehyde-utilizing enzyme, likely reflecting its rapid conversion into downstream metabolites and tight regulation of flux through this branch. In *P. hybrida*, benzaldehyde is detectable but behaves as a highly dynamic intermediate with substantial turnover (Boatright et al., 2004). Flux is likely governed by a combination of factors, including enzyme activity, cofactor availability, and competition with parallel metabolic routes (Moreno-Sánchez et al., 2008; San et al., 2002; Widhalm and Dudareva, 2015). For instance, benzaldehyde can be further oxidized to BA (Long et al., 2009) or reduced to benzyl alcohol (Huang et al., 2022), illustrating how flux may be partitioned across competing enzymatic pathways.

More broadly, biosynthetic enzymes are often organized to facilitate metabolic channeling, enhancing pathway efficiency and minimizing the accumulation of reactive or toxic intermediates (Pareek et al., 2021; Winkel, 2004), a mechanism that may contribute to the efficient turnover of benzaldehyde observed here. In addition, the presence of multiple ADH isoforms, together with the stage-dependent compensation discussed above, further supports a model in which flux can be dynamically adjusted in response to developmental stage or physiological demand. Elucidating how these regulatory layers coordinate pathway output remains an important direction for future studies.

Our cross-species complementation analysis provides additional evolutionary insight into the Phe-derived SA biosynthesis pathway. The ability of *Os*BalSα and *Os*BalSβ to restore SA accumulation in *N. benthamiana* mutants suggests functional conservation at biochemical level (Fig. 2C-D). In *N. benthamiana*, BalS constitutes an essential upstream step required for benzoyl-CoA reduction during pathogen-induced SA biosynthesis. By contrast, rice maintains constitutively high SA levels, and this basal SA production does not depend on BalSα-BalSβ. This discrepancy suggests that rice may deploy redundant or alternative benzoyl-CoA-reducing activities, or channel SA production from benzoyl-CoA through lineage-specific enzymatic routes that bypass BalS.

The conserved requirement for BalR1 in both *N. benthamiana* seedlings and rice (Fig. 3F-G, 5A-B), together with the strong conservation seen in its downstream BEBT-BBO-BSH steps (Liu et al., 2025), points to a modular organization of the Phe-derived pathway, in which upstream nodes exhibit greater lineage-specific plasticity, whereas downstream components form a more conserved core. This modularity likely provides evolutionary flexibility, allowing different plant lineages to diversify their upstream pathway inputs while maintaining robust SA production, particularly during pathogen-induced immune responses.

## Acknowledgements

We would like to thank Dr. Zhonglin Mou (University of Florida) for the Acinetobacter sp. ADPWH-*lux* strain for SA quantification, Dr. Daniel Voytas (University of Minnesota) for the *N. benthamiana* Cas9 transgenic line, Morven Shan (University of British Columbia, UBC) for providing *Marchantia* tissue for gene cloning, and the Arabidopsis Biological Resource Center (ABRC) for the Wiscseq_DsLox375C05 (*Atbalsα*) mutant line. We thank Qingqing Huang (Cornell University) and Conghao Hong (Beijing Forestry University) for technical assistance with structural modeling and site-directed mutagenesis, Dr. Harley Gorden and Dr. Eerik Piirtola (Mass Spectrometry Core Facility at the Wine Research Center of UBC) and Dr. Hsihua Wang (Center of Metabolomics and Proteomics of Sichuan University) for support in metabolite measurement. This work was supported by funds from the National Natural Science Foundation of China (32330008 (Y. Z.), 32300255 (Y.L.)), the Fundamental Research Funds for the Central Universities (YJ202255; YJ202256), and the Canadian Natural Sciences and Engineering Research Council (NSERC) Discovery program (X.L. and Y.Z.), NSERC-CREATE-PRoTECT (X.L.). L.X. is partially supported by a scholarship from China Scholarship Council (CSC).

## Author contributions

X.L. and Y.Z. conceived the study. Y.L. and D.Q. assembled the homolog overexpression constructs, generated rice mutants and performed LC-MS measurement. M.W. contributed to WGS and phylogeny analyses. J.L. and C.L. assisted with confocal microscopy imaging. L.X. conducted the EMS screen and performed all other experiments. L.X. drafted the manuscript. Y.L., X.L. and Y.Z. revised the manuscript. All authors read and approved the final manuscript.

## Materials and Methods

### Plant materials and growth conditions

*N. benthamiana* and rice (*O. sativa*) mutant lines of *BalSα*, *BalSβ* and *BalR1* were generated in this study by CRISPR-Cas9. Rice mutants were generated in the Zhonghua 11 (ZH11) background. *Arabidopsis* Wiscseq T-DNA line Wiscseq_DsLox375C05 (*Atbalsα*) was purchased from the Arabidopsis Biological Resource Center (ABRC) (www.abrc.osu.edu). *N. benthamiana*, rice and *Arabidopsis* plants were grown on soil under long-day conditions (16h-light/8h-dark, 23 ℃/19 ℃ day/night regime) in a growth room with ∼70% relative humidity.

### EMS screen and mapping-by-sequencing

Screening of an ethyl methane sulfonate (EMS)-mutagenized *N. benthamiana* population was performed as previously described (Liu et al., 2025). Briefly, ∼16,000 M2 plants were screened using a modified biosensor-based approach to identify mutants with reduced SA accumulation at 24 hpi with *Pst* DC3000 (OD_600_ = 0.01). For mapping-by-sequencing, the SA-deficient mutant was backcrossed to WT *N. benthamiana* and the F1 plants were self-fertilized. In F₂, individuals exhibiting reduced SA accumulation following *Pst* DC3000 infection were pooled for genomic DNA extraction using a CTAB-based method. Whole-genome sequencing was performed to 30× coverage on an Illumina HiSeq4000 platform with 150-bp paired-end reads (Novogene). Reads were aligned to the reference genome LAB360 (https://apollo.nbenth.com/annotator/index), and single-nucleotide polymorphisms (SNPs) were identified using the Genome Analysis Toolkit (GATK v4). SNP frequency across the genome was plotted to identify linkage region and candidate mutations.

### Mutant generation by CRISPR-Cas9

*N. benthamiana* knockout mutants were generated by Flowering locus T (FT)-mediated CRISPR-Cas9 as described in (Ellison et al., 2021). Genomic sequence of *BalSα*, *BalSβ*, and *BalR1* was analyzed using the VIGS tool (https://vigs.solgenomics.net/) to identify the best target region. 20bp target sequences were selected based on predicted off-target effects and specificity score using Geneious Prime (https://www.geneious.com/). The augmented sgRNA containing the 20bp target sequence and the truncated FT sequence at the 3’ end was amplified from the pEE515 vector using primers listed in Supplementary Table 5. Two or three sgRNAs targeting the same gene were assembled into a spacer multiplexed construct (i.e., TRV2::sgRNA-ABC^BalSα^-FT, TRV2::sgRNA-ABCD^BalSβ^-FT, TRV2::sgRNA-ABC^BalR1^-FT) with AarI restriction enzyme (Thermo Fisher Scientific) and T4 DNA ligase (NEB). The resulting TRV2-sgRNA-FT construct was introduced into *Agrobacterium tumefaciens* strain GV3101. Transformed agrobacteria were grown overnight in LB medium supplemented with appropriate antibiotics at 28 °C with shaking. *Agrobacterium* TRV2 cultures (OD_600_ = 0.8) resuspended in infiltration buffer (10 mM MgCl_2_ and 160 μM acetosyringone) were mixed with TRV1 vector pNJB069 at a 1:1 ratio, incubated at room temperature for 3 h, and infiltrated into the fifth leaf of five-week-old Cas9-overexpressing *N. benthamiana* plants.

*Osbalsα* and *Osbalsβ* knockout mutants were generated using CRISPR-Cas9 as described in Liu et al. (2025). Target sequences were designed with CRISPR-GE (http://skl.scau.edu.cn/), and sgRNA expression cassettes were assembled into the pYLCRISPR/Cas9 binary plasmid via Golden Gate ligation. Rice stable transformants were commercially produced by *Agrobacterium*-mediated transformation of seed-derived green tissues (Boyuan Biotechnology). *Osbalr1* CRISPR knockout mutants were obtained from Biogle Genetech (https://www.biogle.cn). Primers used are provided in Supplementary Table 5.

### Transient gene expression in *N. benthamiana*

A modified pCAMBIA1300 vector (Liu et al., 2025) containing the *Flaveria trinervia* pyruvate orthophosphate dikinase (PDK) intron between the CaMV35S promoter and the *Pisum sativum* trbcS-E9 terminator was used for transient gene silencing. Target sequences of *BEBT* and *ADHs* were PCR-amplified and inserted in both orientations flanking the PDK intron to generate hairpin constructs, using the primers listed in Supplementary Table 5.

For complementation, *NbBalSα* and *NbBalSβ*, or their homologs from *O. sativa*, *Z. mays* and *M. polymorpha* were amplified from genomic DNA or cDNA, and cloned into pCAMBIA1300 vector under the CaMV 35S promoter and trbcS-E9 terminator.

*Agrobacterium* cultures carrying the indicated constructs or empty vector were resuspended in infiltration buffer (10 mM MgCl_2_ and 160 μM acetosyringone) and infiltrated into leaves of *N. benthamiana* plants (OD_600_ = 0.4). Three days later, leaves were challenged with *Pst* DC3000 (OD_600_ = 0.02), and SA levels were quantified 8 h post inoculation.

### Measurement of SA and SAG

To quantify pathogen-induced SA, SAG and SGE levels, five-week-old *N. benthamiana* plants were syringe-infiltrated with *Pst* DC3000. Bacteria were grown overnight, pelleted, and resuspended in 10 mM MgCl_2_ to the desired densities, and infiltrated into leaves using a needleless syringe. For HPLC-based measurements, plants were infiltrated at OD_600_ = 0.02 and sampled at 8 hpi. For large-scale analysis using the biosensor assay, plants were infiltrated at OD_600_ = 0.01 and sampled at 24 hpi.

For rapid SA quantification, a biosensor-based assay using *Acinetobacter* sp. ADPWH_*lux* was performed as described by Liu et al. (2025). Briefly, two leaf discs (7 mm in diameter) were collected from the infiltrated area, placed into PCR plates, and boiled in 90 μL LB at 95 °C for 10 min. After cooling, 50 μL of the extract was transferred into a white 96-well plate and mixed with 50 μL of freshly prepared biosensor culture (OD_600_ = 0.4). After incubation at 37 °C for 1 h, luminescence signals were measured using a microplate reader. SA concentrations were determined using a standard curve generated from pure SA standards processed in parallel with the samples.

For HPLC-based measurement, SA quantification was performed as described by Tian et al. (2021). ∼100 mg of leaf tissue was harvested (four biological replicates per genotype), flash frozen in liquid nitrogen, and finely ground using TissueLyser II (QIAGEN). SA was extracted sequentially with 90% and 100% methanol, 5% (w/v) trichloroacetic acid and extraction buffer (ethylacetate acid/cyclopentane/isoporopanol, 100:99:1, v/v), and the final residue was dissolved in 500 μL mobile phase (0.2 M potassium acetate, 0.5 mM EDTA, pH 5). 10 μL of supernatant was injected into an HPLC system equipped with a fluorescence detector (Ex 295 nm, Em 405 nm; Shimadzu) and an XBridge C18 column (4.6 × 100 mm, 3.5 μm; Waters, Cat. 186003033). An isocratic flow (1.3 mL/min) was applied with a total run time of 10 min. Quantitation was performed using a standard curve generated with pure SA standards. Data were integrated and processed using LabSolution software (Shimadzu, v5.111).

SAG quantification was performed using LC–MS following Liu et al. (2025). Briefly, ground plant tissue was extracted sequentially with 80% and 100% methanol, and 2 μl of the clarified extract was analyzed by LC–MS/MS (Nexera UHPLC LC-30A coupled to an AB SCIEX Triple Quad 5500). SA and SAG were identified and quantified using pure chemical standards. Metabolites were separated on a Hypersil Gold C18 column (100 × 2.1 mm, 1.9 μm) at 40 °C with a 0.4 ml min⁻¹ flow rate using a water (0.1% formic acid)/acetonitrile gradient. Detection was carried out in negative-mode ESI using MRM, and quantification was based on standard curves for generated from pure SA and SAG standards.

### Measurement of benzaldehyde by GC-QqQ-MS/MS

Metabolite extraction was performed as described in Liu et al. (2025). Briefly, ∼100 mg of leaf tissue collected at 8 hpi with *Pst* DC3000 (OD_600_ = 0.02) from 5-week-old *NbBEBT*-silenced WT, *balsα* and *balsβ* plants was ground in liquid nitrogen and extracted in 500 μL hexane. Samples were sonicated for 5 min and rotated overnight at 4 °C, then centrifuged at 3,300 g for 15 min. The organic phase was analyzed using an Agilent 7890 GC system coupled to Agilent 7000A triple quad and Agilent DB-1701 column (122-0732; 30 m × 0.25 mm × 0.25 mm; J&W Scientific). Helium was used as carrier gas (1.1 mL/min) in pulsed splitless mode. The oven program started at 45 °C (2 min hold), ramped to 240 °C at 10 °C/min, then to 280 °C at 35 °C/min (5 min hold), with a total running time of 27.64 min and solvent delay of 3.5 min. Injector, ion source, and quadrupole temperatures were 250, 230, and 150 °C, respectively. Electron impact ionization (70 eV) was used, and compounds were detected in MRM mode with 20 eV collision energy, using He (2.25 mL/min) and N₂ (1.5 mL/min) as quenching and collision gases, respectively. Benzaldehyde was identified by retention time and characteristic transition (m/z 106 > 77) relative to pure chemical standard (benzaldehyde, 09143, Sigma-Aldrich). Quantification was based on calibration curve using Agilent MassHunter QQQ Quantitative Analysis (v10.2).

### Protein purification and *in vitro* enzymatic assay

Protein purification and activity assays were performed as described previously (Huang et al., 2022; Liu et al., 2025) with minor modifications. The coding sequences of *NbBalSα, NbBalSβ, NbBalR1* and codon-optimized *OsBalR1* were amplified and cloned into a modified pMAL-c2x vector containing an N-terminal His-MBP tag. The codon-optimized *OsBalR1* was commercially synthesized (Thermo Fisher Scientific). Constructs were transformed into *E. coli* Rosetta cells. Cultures were grown at 37 °C to OD₆₀₀ ≈ 0.5, induced with 0.5 mM IPTG, and incubated at 28 °C for 3 h (*Nb*BalSα and *Nb*BalSβ) or 18 °C for 18 h (*Nb*BalR1 and *Os*BalR1). Cells were harvested, washed, and lysed by sonication in lysis buffer (50 mM Tris pH 7.5, 500 mM NaCl, 10% glycerol, 0.2% Triton X-100, 1 mM PMSF, 1 mM DTT). Lysates were clarified by centrifugation (15,000 rpm, 4 °C, 40 min), and the supernatants were loaded onto Ni-NTA agarose (QIAGEN) for affinity purification. Eluted proteins were dialyzed and verified by SDS-PAGE.

For *Nb*BalS assays, 100 μL reaction contained 1.3 μg purified *Nb*BalSα and *Nb*BalSβ proteins (1:1 molar ratio), 50 mM Bis-Tris buffer (pH 6.5), 200 μM benzoyl-CoA (Sigma-Aldrich), and 2 mM NADPH (Sigma-Aldrich). Reactions were incubated at 28 °C for 30 min and terminated with 100% ice-cold methanol. For kinetic analyses, benzoyl-CoA concentrations ranging from 80 μM to 1.6 mM were tested in the presence of 4 mM NADPH.

For *Nb*BalR1 assays, 100 µL reactions were conducted at 28 °C in 50 mM Tris-HCl buffer (pH 7.5) containing 2 μg purified *Nb*BalR1, 2 mM benzaldehyde (Sigma-Aldrich), and 2 mM NADH or NADPH (Sigma-Aldrich). To determine the *K*_m_ for benzaldehyde, substrate concentrations from 2 µM to 100 µM were tested in the presence of 2 mM NADPH or NADH. To determine the *K*_m_ for NADPH, cofactor concentrations ranging from 10 µM to 1 mM were tested in the presence of 150 µM benzaldehyde. For *Os*BalR1 assays, benzaldehyde concentrations ranging from 50 µM to 1500 µM were tested in the presence of 2 mM NADPH, while NADPH concentrations ranging from 10 µM to 1 mM were tested in the presence of 1 mM benzaldehyde. Reactions were incubated for 30 min and stopped with 100% ice-cold methanol. Product formation was quantified by HPLC-DAD. *K*_m_ and V_max_ values were obtained by nonlinear regression fitting to the Michaelis-Menten equation using GraphPad Prism (v10.2.0) based on triplicate measurements.

### HPLC analysis of reaction products

Reaction products were analyzed using an Agilent 1260 Infinity II HPLC system equipped with a diode array detector (DAD) and an InfinityLab Poroshell HPH-C18 column (4.6 × 100 mm, 2.7 μm; Agilent). The mobile phases were 0.13% trifluoroacetic acid in water (A) and 0.1% trifluoroacetic acid in acetonitrile (B), at a flow rate of 1 mL/min and a column temperature of 30 °C. 10 μL of the product was injected and separated using a 25 min linear gradient: 10-90% B over 15 min, held for 5 min, and re-equilibrated back to 10% B. Benzaldehyde and benzyl alcohol were detected at 248 nm and 210 nm, respectively, and identified by comparison with authentic standards. Quantification was based on standard curves using Agilent OpenLab CDS analysis software (v2.6).

### Phylogenetic analysis

Homologs of *Nb*BalSα and *Nb*BalSβ were identified from 83 representative plant species using the corresponding *N. benthamiana* protein sequences as BLASTp queries against publicly available protein databases (Supplementary Table 2). Putative homologs were retained only if they passed reciprocal BLASTp validation, where the *Nb*BalSα or *Nb*BalSβ was recovered as the top hit when queried against the *N. benthamiana* database. Reciprocally validated sequences were further examined for conserved domain architecture using InterProScan (https://www.ebi.ac.uk/interpro/), and only those containing the characteristic SDR-related domains were included in phylogenetic analyses. Protein sequences were aligned using MAFFT (v7.490) with default parameters, followed by manual inspection and trimming to remove poorly aligned terminal regions. Maximum-likelihood phylogenetic trees were inferred with IQ-TREE (v2.2.2.6) using the best-fit model selected based on BIC scores with 1,000 ultrafast bootstrap replicates. Trees were visualized and annotated using FigTree (v1.4.4), with bootstrap support values (%) displayed at corresponding nodes.

### Protein structure analysis

Tertiary structures of BalSα, BalSβ, and their homologs were predicted using AlphaFold2 (https://colab.research.google.com/github/sokrypton/ColabFold/blob/main/AlphaFold2.ipynb). Predicted structures were visualized and Pairwise protein alignments were performed in PyMOL (v3.1.0) to assess structural conservation. Interfacial contacts between BalSα and BalSβ were further analyzed using RING (v3.0) with default parameters and input model generated by AlphaFold. Predicted hydrogen bonds, salt bridges, π–π interactions, and van der Waals contacts were mapped onto the structural model for identification of key interfacial residues.

### Site-directed mutagenesis

Point mutations (Y257A and E264A/Y265A) were introduced into *BalSα* using PCR-based site-directed mutagenesis with primers listed in Supplementary Table 5. Mutant constructs were verified by Sanger sequencing. Expression and purification of protein variants (BalSα^Y257A^ or BalSα^E264A/Y265A^) were performed as described above for WT BalSα.

### *In vitro* pull-down assay

For protein-protein interaction assays, *BalSβ* was cloned into the pET-24c vector without an N-terminal His-MBP tag, whereas *BalSα* and its mutant variants were cloned into the modified pMAL-c2x vector described earlier. Constructs were expressed individually in *E. coli* Rosetta cells. After IPTG induction and cell lysis by sonication, the soluble fractions were applied to Ni-NTA resin (Qiagen), washed extensively, and bound proteins were eluted with 250 mM imidazole. Eluted fractions were analyzed by SDS-PAGE followed by Coomassie Brilliant Blue staining to assess the interaction between BalSα or its variants and BalSβ.

### Subcellular localization of GFP-tagged proteins

To generate GFP fusion constructs, the CDSs of *NbBalSα*, *NbBalSβ*, and *NbBalR1* were amplified and fused in-frame to the C terminus of EGFP in the pCAMBIA1300 vector. For BiFC constructs, the CDSs of *NbBalSα* and *NbBalSβ* were fused in-frame to the C terminus of nEYFP and cEYFP, respectively, in the pCAMBIA1300 vector. The peroxisomal marker mCherry-SKL was generated by fusing the peroxisomal targeting signal SKL (Ser-Lys-Leu) to the C terminus of mCherry in the pCAMBIA1300 vector. *Agrobacterium* strains (OD_600_ = 0.6) carrying fluorescent fusion constructs (i.e., EGFP-BalSα, EGFP-BalSβ, EGFP-BalR1, nEYFP-BalSα, and cEYFP-0BalSβ) were mixed with strains carrying mCherry-SKL at a 1:1 ratio and co-infiltrated into five-week-old *N. benthamiana* leaves. At 48 hpi, fluorescence signals were examined in abaxial epidermal cells. Images were acquired using an Evident IXplore SpinSR Spinning Disk Confocal Microscope (Olympus) equipped with a Yokogawa CSU-W1 unit. GFP was excited using a 488 nm laser line and detected with a 525/50 nm emission filter, YFP was excited using a 514 nm laser line and detected with a 535/50 nm emission filter, and mCherry was excited using a 561 nm laser line and detected with a 617/73 nm emission filter.

### Gene expression analysis

RNA extraction and quantitative real-time PCR (qRT-PCR) were performed as previously described (Xu et al., 2026). Briefly, ∼100 mg of leaf tissue was harvested at 6 hpi with *Pst* DC3000 (four biological replicates per genotype). Total RNA was isolated using the EZ-10 Spin Column Plant RNA Mini-Preps Kit (Bio Basic) and reverse transcribed into cDNA using the OneScript Reverse Transcriptase kit (ABM). qRT-PCR was performed with SYBR Premix Ex Taq II kit (TAKARA) using primers listed in Supplementary Table 5. *NbPP2A* was used as the internal reference gene.

### Statistical analysis

Error bars represent standard deviations from three to six biological replicates. Statistical analyses were performed using GraphPad Prism 10 (v10.2.0). Significance was determined using two-tailed Student’s *t*-test or one-way ANOVA with Tukey’s post hoc test, as indicated in figure legends.

## Supplementary figure legends

**Supplementary Figure S1. Identification and characterization of low SA mutants in *N. benthamiana*.**

**A)** Free SA levels in five-week-old WT and three SA-deficient mutants at 24 hpi with *Agrobacterium* (OD_600_ = 0.4) carrying either empty vector or TIR gene *At2g32140*, as measured using HPLC.

**B)** Free SA levels in five-week-old F_1_ hybrids from crosses between WT and mutant lines 4-66, 13-270, and 15-328 at 24 hpi with *Pst* DC3000 (OD_600_ = 0.01), as measured by a biosensor-based method.

**C)** Free SA levels in four-week-old F_2_ segregants derived from the same crosses as in (**B**) at 24 hpi with *Pst* DC3000 (OD_600_ = 0.01) measured by a biosensor-based method. Induced SA levels in WT and SA-deficient mutants grown are indicated by blue and green lines, respectively. The red line represents the expected 1:3 segregation ratio, calculated from the total number of F₂ plants measured, which is consistent the observed segregation pattern of SA levels. The number of F₂ plants used for DNA extraction and WGS is noted alongside.

**D)** Sanger sequencing chromatograms of two independent CRISPR mutants of *BalSα*. Line *balsα #3* carries a 7-bp deletion at sgRNA target site A, a 146-bp deletion spanning target sites B and C, and a G-to-T substitution at target site C. Line *balsα #6* contains a 246-bp deletion spanning target sites A and B.

**E)** Morphology of five-week-old SA-deficient mutants (4-66 and 15-328), *balsα* CRISPR mutants, and WT. Scale bar, 1 cm.

Error bars represent mean ± s.d. ns, no significant difference. Letters indicate statistically significant differences (one-way ANOVA with Tukey’s test, n = 4 independent plants).

**Supplementary Figure S2. Genotypic validation and morphological characterization of *balsβ* CRISPR mutants.**

**A)** Sanger sequencing chromatograms of two independent CRISPR mutants of *BalSβ*. Line *balsβ #1* carries a 100-bp deletion spanning sgRNA target sites A and B. Line *balsβ #9* contains a 1893-bp deletion spanning target sites B and C.

**B)** Morphology of five-week-old SA-deficient mutant 13-270, *balsβ* CRISPR mutants, and WT. Scale bar, 1 cm.

**Supplementary Figure S3. Protein purification and enzymatic activity of BalSα and BalSβ.**

**A)** Coomassie stained SDS-PAGE gel showing purified BalSα and BalSβ, and the His-MBP tag expressed in *E. coli*. Total, bacterial cultures with (+) or without (-) IPTG induction; WCL, whole cell lysate; Eluted, fractions after Ni-NTA affinity purification; Dialyzed, fractions after dialysis; M, marker. Black triangle denotes His-MBP-tagged BalSα (∼74 kDa) and BalSβ (∼73 kDa), while blue triangle denotes the His-MBP tag.

**B)** Estimation of protein concentration using Bovine serum albumin (BSA) standards.

**C)** Kinetic curves of BalSα-BalSβ reaction with varying concentrations of benzoyl-CoA. Data from three replicates were fitted using non-linear Hill regression in Prism 10 (v10.2.0).

**Supplementary Figure S4. Subcellular localization of EGFP-BalS subunits to the peroxisome.**

**A-C)** EGFP-BalSα (**A**), EGFP-BalSβ (**B**), or N-terminal fragment of EYFP (nEYFP)-BalSα together with C-terminal fragment of EYFP (cEYFP)-BalSβ (**C**) were transiently co-expressed with the peroxisome marker mCherry-PTS1 in five-week-old *N. benthamiana* leaves and visualized by confocal microscopy. EGFP/EYFP fluorescence, mCherry fluorescence (peroxisome), and merged images are shown. Images were acquired at 40× and 60× magnification.

**Supplementary Figure S5. Presence, evolutionary distribution, and structural conservation of BalSα and BalSβ homologs across representative plant species.**

Protein structures of selected BalSα and BalSβ homologs were predicted using AlphaFold2 and superimposed onto the corresponding *Nb*BalSα or *Nb*BalSβ using pyMOL, with the RMSD values indicated. Filled green circles and open circles denote the presence and absence of homologs, respectively, with the number showing the copy number of homologs identified in each species. Stars mark inferred gene gain events. The phylogenetic framework is based on the accepted species tree topology from TimeTree (https://timetree.org/), with branch lengths not proportional to primary sequence divergence. For clarity, some eudicot and monocot species are omitted due to space limitations.

**Supplementary Figure S6. Phylogeny of *Nb*BalSα and its homologs in plants.**

Phylogenetic tree of BalSα homologs was constructed using protein sequences retrieved from the BLASTp searches across 84 representative plant species spanning major Archaeplastida lineages. Sequences were aligned using MAFFT (v7.490), and the tree was built using IQ-TREE (v2.2.2.6) under the maximum-likelihood method with 1,000 ultrafast bootstrap replicates. The resulting tree was visualized and annotated in FigTree (v1.4.4). Ultrafast bootstrap support values (%) are shown next to nodes. Reported genes encoding BalSα are highlighted in bold. B.A., basal angiosperms.

**Supplementary Figure S7. Phylogeny of *Nb*BalSβ and its homologs in plants.**

Phylogenetic tree of BalSβ homologs was constructed using protein sequences retrieved from the BLASTp searches across 84 representative plant species spanning major Archaeplastida lineages. Sequences were aligned using MAFFT (v7.490), and the tree was built using IQ-TREE (v2.2.2.6) under the maximum-likelihood method with 1,000 ultrafast bootstrap replicates. The resulting tree was visualized and annotated in FigTree (v1.4.4). Ultrafast bootstrap support values (%) are shown next to nodes. Reported genes encoding BalSβ are highlighted in bold. B.A., basal angiosperms.

**Supplementary Figure S8. Validation of rice CRISPR lines by Sanger sequencing and analysis of Arabidopsis *balsβ* mutants.**

**A)** Sanger sequencing chromatograms of two independent mutants of each gene, *OsBalSα* and *OsBalSβ*. Line *Osbalsα #1* carries a 5-bp deletion at sgRNA target site A and a 1-bp insertion at site B. *Osbalsα #2* has a substitution/insertion replacing CGA with AAAAAC at site A. *Osbalsβ #1* and *Osbalsβ #2* harbor 201-bp and 203-bp deletions between sites A and B, respectively.

**B)** Structure of the single *AtBalSβ* gene showing location of the T-DNA insertion in the mutant line *Atbalsβ*. Exons are represented by green rectangles, UTRs by grey rectangles, and introns by connecting black lines. The primers used for PCR genotyping are depicted with blue arrows.

**C)** Morphology of three-week-old Col-0, *sid2* (mutating *ICS1*) and *Atbalsβ* plants grown under long-day conditions. Scale bar, 1 cm.

**D)** Free SA levels of four-week-old soil-grown *Arabidopsis* plants of the indicated genotypes at 16 hpi with *Pst* DC3000 avrRpt2 (OD_600_ = 0.005) or MgCl_2_, as measured by HPLC. Asterisks indicate statistically significant differences (two-tailed Student’s *t*-tests, \**P* < 0.05, n = 4 independent plants). ns, no significant difference.

**Supplementary Figure S9. Structural analysis of the BalSα-BalSβ interface.**

**A-C)** Ribbon diagrams of the BalSα-BalSβ heterodimer highlighting predicted interface residues. Tyr257 in BalSα was predicted to simultaneously form a hydrogen bond with Thr232 and engage in π-π stacking with Tyr235 of BalSβ, whereas Glu264 and Tyr265 were predicted to form inter-chain hydrogen bonds with Lys227 and Gly253 of BalSβ, respectively. Zoom-in views show BalSα residues Tyr257 (**A**), Glu264 (**B**), and Tyr265 (**C**) (green sticks) and their interacting residues in BalSβ (orange sticks), with distances indicated. Hydrogen bonds (blue dashed lines) and π-π stacking interactions (orange dashed lines) were identified using RING.

**D)** Coomassie-stained SDS-PAGE gel from *in vitro* pull-down assays showing co-purification of untagged BalSβ with 6×His-MBP-tagged BalSα, its mutant variants or the tag alone using Ni-NTA resin. Proteins were expressed in *E. coli*, equal amounts of lysates were incubated with the same amount of BalSβ and Ni-NTA beads, and eluates were analyzed by SDS-PAGE. Filled triangle indicates BalSβ (∼29 kDa), open triangle indicates His-MBP-BalSα and its variants (∼74 kDa), and blue triangle indicates the His-MBP control (∼43 kDa).

**E)** *In vitro* benzoyl-CoA reduction into benzaldehyde by BalSβ together with BalSα, BalSα^Y257A^ or BalSα^E264A/Y265A^. Reactions containing BalSβ and BalSα or its variants were incubated with benzyl-CoA and NADPH at 28 °C for 30 min. Benzaldehyde formation was verified by HPLC-DAD against an authentic pure standard.

**F)** Free SA levels in *balsα* mutants transiently overexpressed with empty vector (EV), BalSα, BalSα^Y257A^ or BalSα^E264A/Y265A^ at 8 hpi with *Pst* DC3000 (OD_600_ = 0.02), as measured by a biosensor-based method. *N. benthamiana* leaves were infiltrated with *Agrobacterium* carrying the overexpression constructs (OD_600_ = 0.4) three days before *Pst* DC3000 inoculation. Error bars represent mean ± s.d. (one-way ANOVA with Tukey’s test, n = 4 independent plants). Letters indicate statistically significant differences.

**G-H)** Sequence alignment of regions surrounding key interface residues in BalSα (**G**) and BalSβ

(**H**) across representative species. Identical residues are highlighted in blue.

**I)** Ribbon diagram of BalS heterodimer and superimposition of the two subunits showing high structural symmetry. BalSα (yellow) and BalSβ (blue) exhibit strong structural overlap (RMSD = 0.737 Å). Secondary structural elements are colored as follows: α-helices (red), β-sheets (yellow), and loops (green). Insets show top, front, and back views of the superimposed subunits. Models were generated with AlphaFold2 and visualized in PyMOL.

**J)** Sequence alignment of BalSα and BalSβ. Predicted secondary structure elements based on AlphaFold are shown above the alignment (α-helices in red and β-sheets in yellow). Conserved residues are indicated by grey shading.

**Supplementary Figure S10. Functional characterization and regulation of *Nb*BalR1 in SA biosynthesis.**

**A)** Coomassie-stained SDS-PAGE gel showing purification of His-MBP-tagged BalR1 expressed in *E. coli*. Total, bacterial cultures with (+) or without (-) IPTG induction; WCL, whole cell lysate; S, soluble lysate after sonication; Post-binding, flow-through after Ni-NTA binding; Post-washing, flow-through after washing; Dialyzed: fractions after dialysis. M, marker. Black triangle indicates purified His-MBP tagged BalR1 (∼82 kDa). BSA standards were used to estimate protein concentration.

**B)** Kinetic analysis of BalR1 activity with varying concentrations of NADH in the presence of benzaldehyde. Data from three replicates were fitted using non-linear Hill regression in Prism 10 (v10.2.0).

**C-E, G)** Relative expression of *NbBalR1* (**C**), *NbBalSα* (**D**), and *NbBalSβ* (**E**) in the indicated genotypes at 6 hpi with *Pst* DC3000 (OD_600_ = 0.02), and of *ADH* genes in hpRNA-silenced plants relative to empty vector (EV) controls (**G**), measured by qRT–PCR (log₂ scale). Error bars represent mean ± s.d. ns, no significant difference. Letters indicate statistically significant differences (one-way ANOVA with Tukey’s test, n = 4 independent plants). Asterisks indicate statistically significant differences (two-tailed Student’s *t*-tests, \**P* < 0.05, n = 4 independent plants).

**F)** Sanger sequencing chromatograms of two independent CRISPR mutants of *NbBalR1*. Line *balr1 #1* carries an 817-bp deletion between sgRNA target site A and B. Line *balr1 #2* carries a 1-bp insertion at site A.

**Supplementary Figure S11. Validation of rice CRISPR lines by Sanger sequencing and purification of *Os*BalR1.**

**A)** Sanger sequencing chromatograms of two independent CRISPR mutants of *OsBalR1*. Line *Osbalr1 #1* and line *Osbalr1 #2* carry 1-bp and 6-bp deletions at target site A, respectively.

**B)** Coomassie-stained SDS-PAGE gel showing the purified *Os*BalR1 expressed in *E. coli*. Total, bacterial cultures with (+) or without (-) IPTG induction; WCL, whole cell lysate; IS, insoluble fraction; S, soluble lysate after sonication; Post-binding, flow-through after Ni-NTA binding; Post-washing, flow-through after washing; Eluted, eluate; Dialyzed: fractions after dialysis. M, marker. Black triangle indicates purified His-MBP tagged *Os*BalR1 (∼82 kDa). BSA standards were used to estimate protein concentration.

**Supplementary Table 1. Candidate genes identified by whole-genome sequencing (WGS).**

**Supplementary Table 2. Sources and database versions used for phylogenetic analysis.**

**Supplementary Table 3. Expression of annotated dehydrogenases among XopQ-responsive transcripts.**

**Supplementary Table 4. The top 50 co-expressed genes of *OsBalR1* (*Os04g15920.1*).**

**Supplementary Table 5. Primers used in this study.**

