## Supplemental Figures for "Identification of Reductases Catalyzing Benzyl Alcohol Formation during Salicylic Acid Biosynthesis in Plants"

#### Supplementary Figure S1

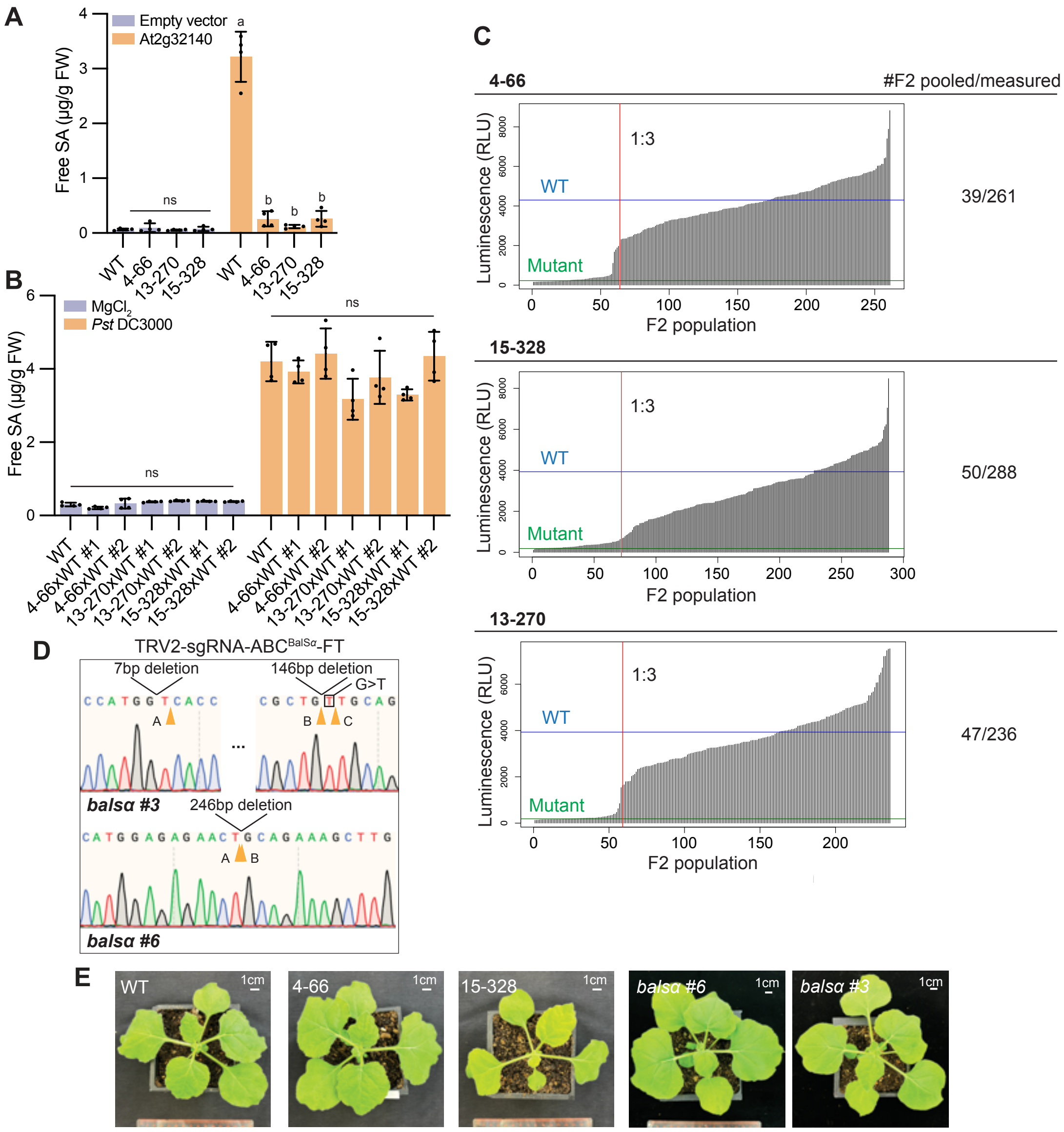


**Supplementary Figure S1. Identification and characterization of low SA mutants in *N. benthamiana*.**
**A)** Free SA levels in five-week-old WT and three SA-deficient mutants at 24 hpi with *Agrobacterium* ($OD_{600}$ = 0.4) carrying either empty vector or TIR gene *At2g32140*, as measured using HPLC.
**B)** Free SA levels in five-week-old F1 hybrids from crosses between WT and mutant lines 4-66, 13-270, and 15-328 at 24 hpi with *Pst* DC3000 ($OD_{600}$ = 0.01), as measured by a biosensor-based method.
**C)** Free SA levels in four-week-old $F_2$ segregants derived from the same crosses as in (**B**) at 24 hpi with *Pst* DC3000 ($OD_{600}$ = 0.01) measured by a biosensor-based method. Induced SA levels in WT and SA-deficient mutants grown are indicated by blue and green lines, respectively. The red line represents the expected 1:3 segregation ratio, calculated from the total number of $F_2$ plants measured, which is consistent the observed segregation pattern of SA levels. The number of $F_2$ plants used for DNA extraction and WGS is noted alongside.
**D)** Sanger sequencing chromatograms of two independent CRISPR mutants of *BalSα*. Line *balsα #3* carries a 7-bp deletion at sgRNA target site A, a 146-bp deletion spanning target sites B and C, and a G-to-T substitution at target site C. Line *balsα #6* contains a 246-bp deletion spanning target sites A and B.
**E)** Morphology of five-week-old SA-deficient mutants (4-66 and 15-328), *balsα* CRISPR mutants, and WT. Scale bar, 1 cm.
Error bars represent mean ± s.d. ns, no significant difference. Letters indicate statistically significant differences (one-way ANOVA with Tukey's test, n = 4 independent plants).

### Supplementary Figure S2

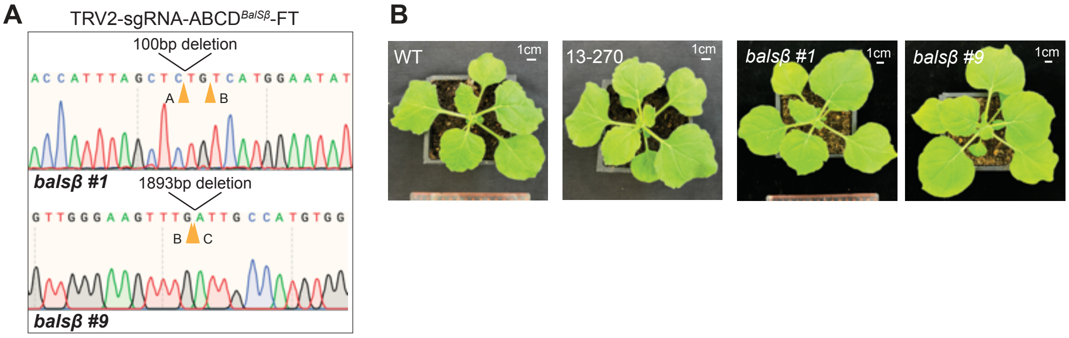


**Supplementary Figure S2. Genotypic validation and morphological characterization of *balsβ* CRISPR mutants.**
**A)** Sanger sequencing chromatograms of two independent CRISPR mutants of *BalSβ*. Line *balsβ #1* carries a 100-bp deletion spanning sgRNA target sites A and B. Line *balsβ #9* contains a 1893-bp deletion spanning target sites B and C.
**B)** Morphology of five-week-old SA-deficient mutant 13-270, *balsβ* CRISPR mutants, and WT. Scale bar, 1 cm.

#### Supplementary Figure S3

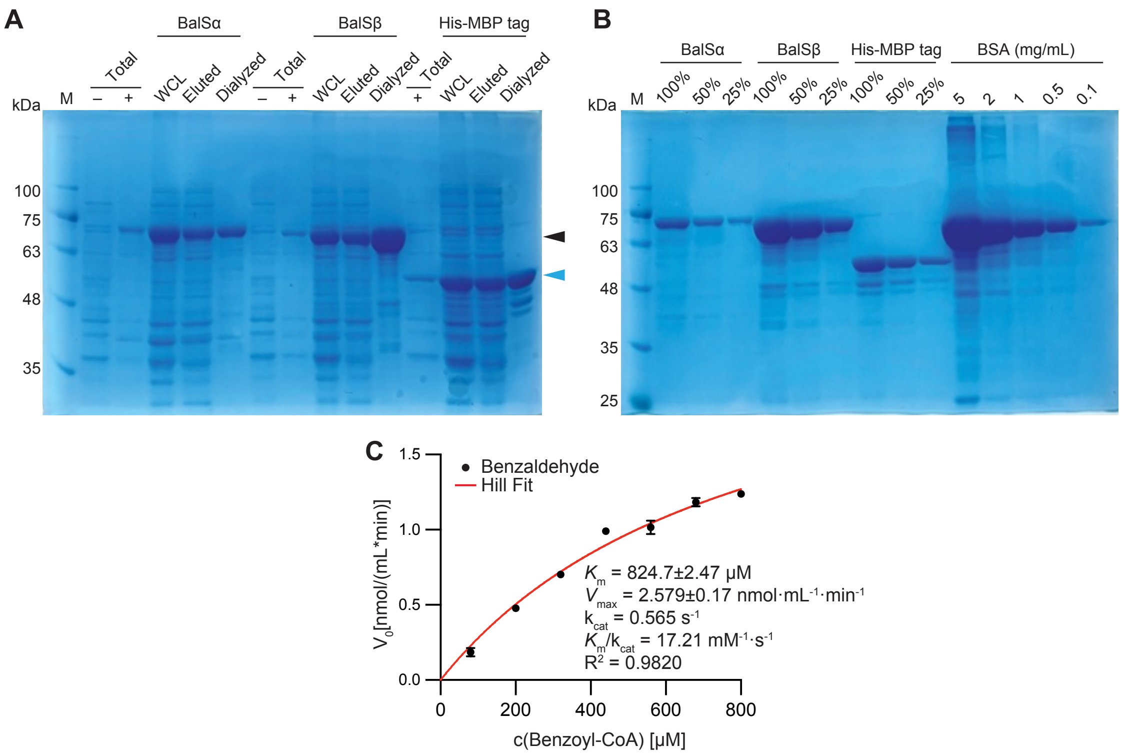


**Supplementary Figure S3. Protein purification and enzymatic activity of BalSα and BalSβ.**

**A)** Coomassie stained SDS-PAGE gel showing purified BalSα and BalSβ, and the His-MBP tag expressed in *E. coli*. Total, bacterial cultures with (+) or without (-) IPTG induction; WCL, whole cell lysate; Eluted, fractions after Ni-NTA affinity purification; Dialyzed, fractions after dialysis; M, marker. Black triangle denotes His-MBP-tagged BalSα (~74 kDa) and BalSβ (~73 kDa), while blue triangle denotes the His-MBP tag.

**B)** Estimation of protein concentration using Bovine serum albumin (BSA) standards.

**C)** Kinetic curves of BalSα-BalSβ reaction with varying concentrations of benzoyl-CoA. Data from three replicates were fitted using non-linear Hill regression in Prism 10 (v10.2.0).

### Supplementary Figure S4

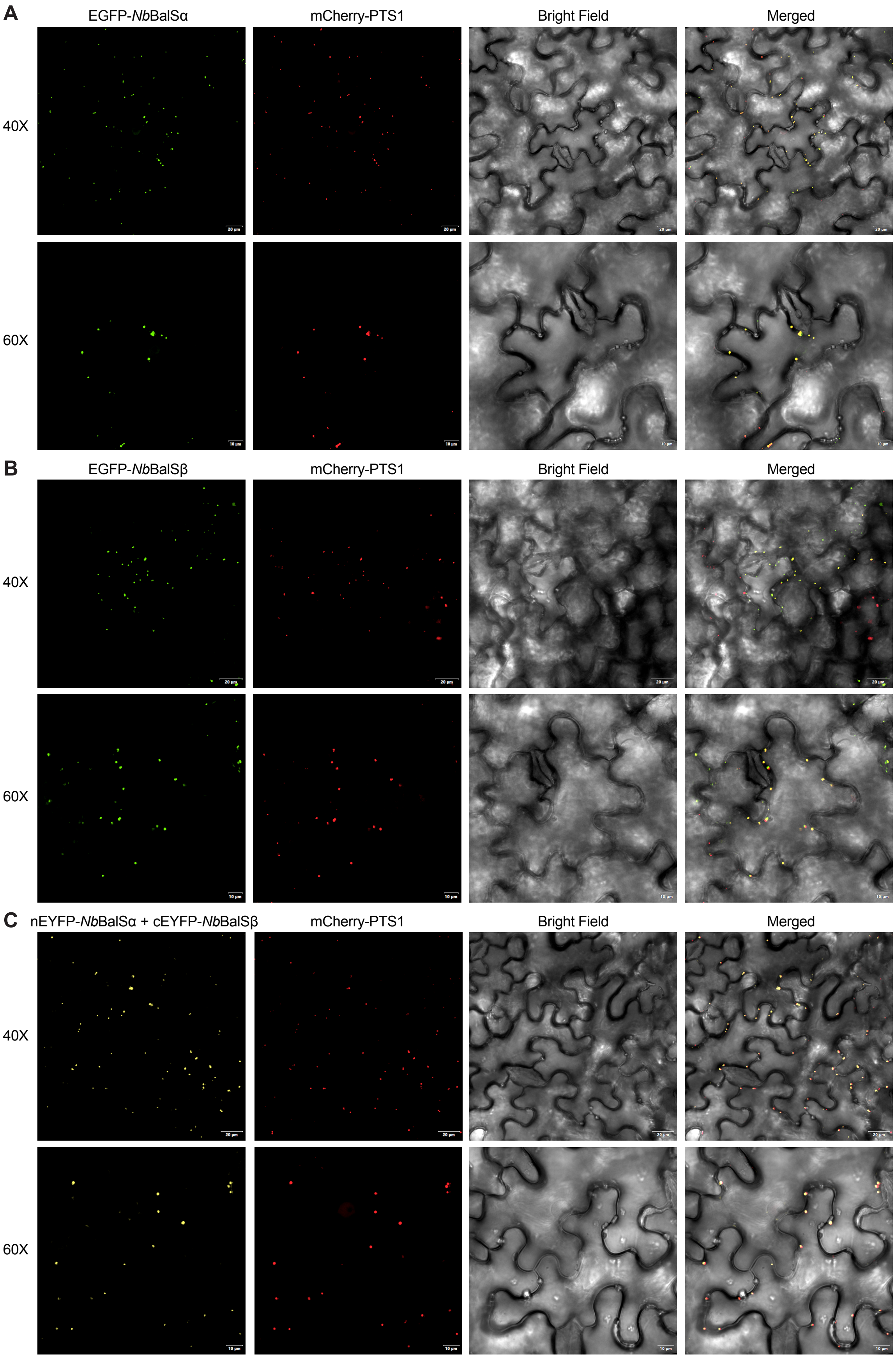


**Supplementary Figure S4**. **Subcellular localization of EGFP-BalS subunits to the peroxisome.**
**A-C)** EGFP-BalSα (**A**), EGFP-BalSβ (**B**), or N-terminal fragment of EYFP (nEYFP)-BalSα together with C-terminal fragment of EYFP (cEYFP)-BalSβ (**C**) were transiently co-expressed with the peroxisome marker mCherry-PTS1 in five-week-old *N. benthamiana* leaves and visualized by confocal microscopy. EGFP/EYFP fluorescence, mCherry fluorescence (peroxisome), and merged images are shown. Images were acquired at 40× and 60× magnification.

#### Supplementary Figure S5

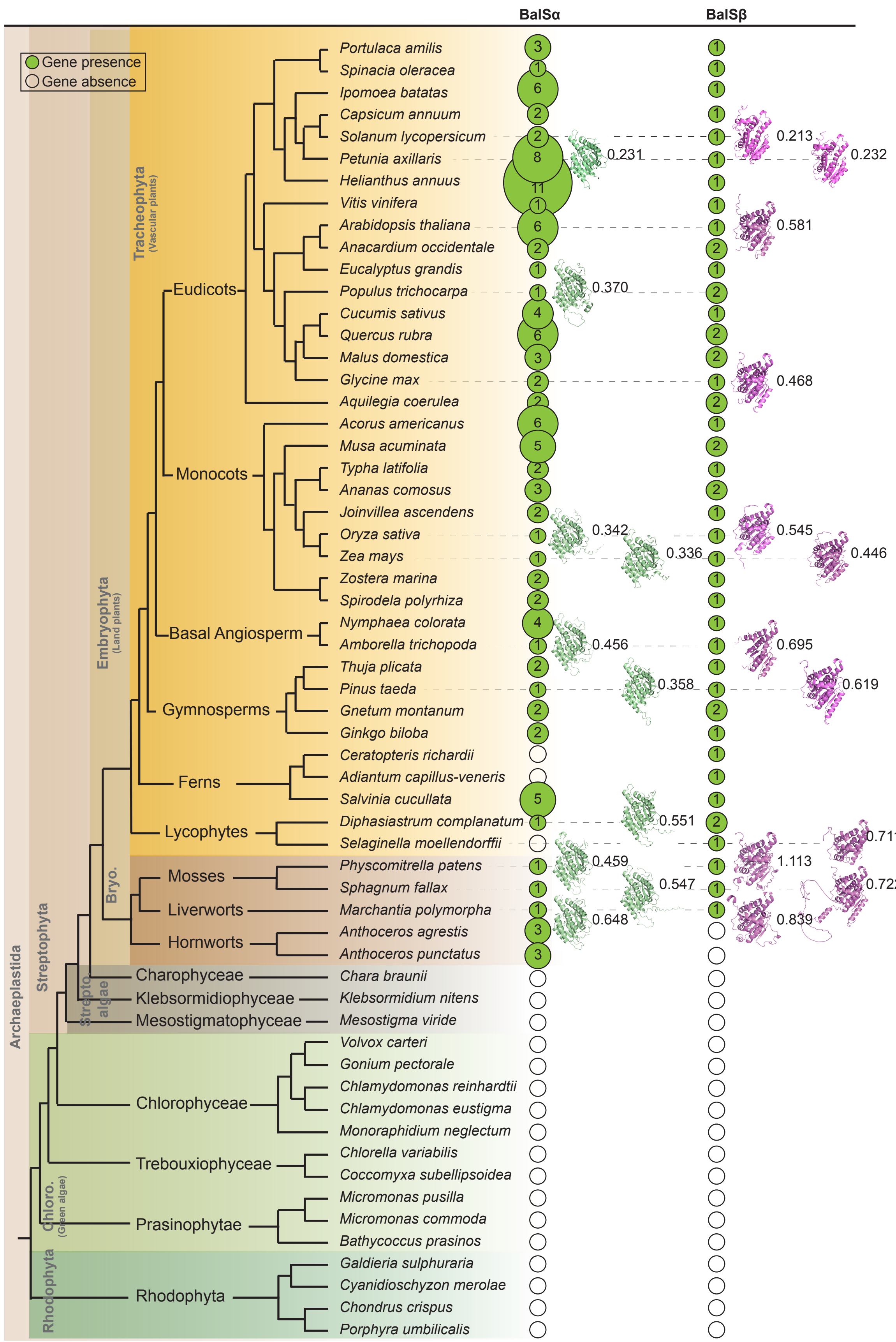


**Supplementary Figure S5. Presence, evolutionary distribution, and structural conservation of BalSα and BalSβ homologs across representative plant species.**
Protein structures of selected BalSα and BalSβ homologs were predicted using AlphaFold2 and superimposed onto the corresponding *Nb*BalSα or *Nb*BalSβ using pyMOL, with the RMSD values indicated. Filled green circles and open circles denote the presence and absence of homologs, respectively, with the number showing the copy number of homologs identified in each species. Stars mark inferred gene gain events. The phylogenetic framework is based on the accepted species tree topology from TimeTree (https://timetree.org/), with branch lengths not proportional to primary sequence divergence. For clarity, some eudicot and monocot species are omitted due to space limitations.

### Supplementary Figure S6

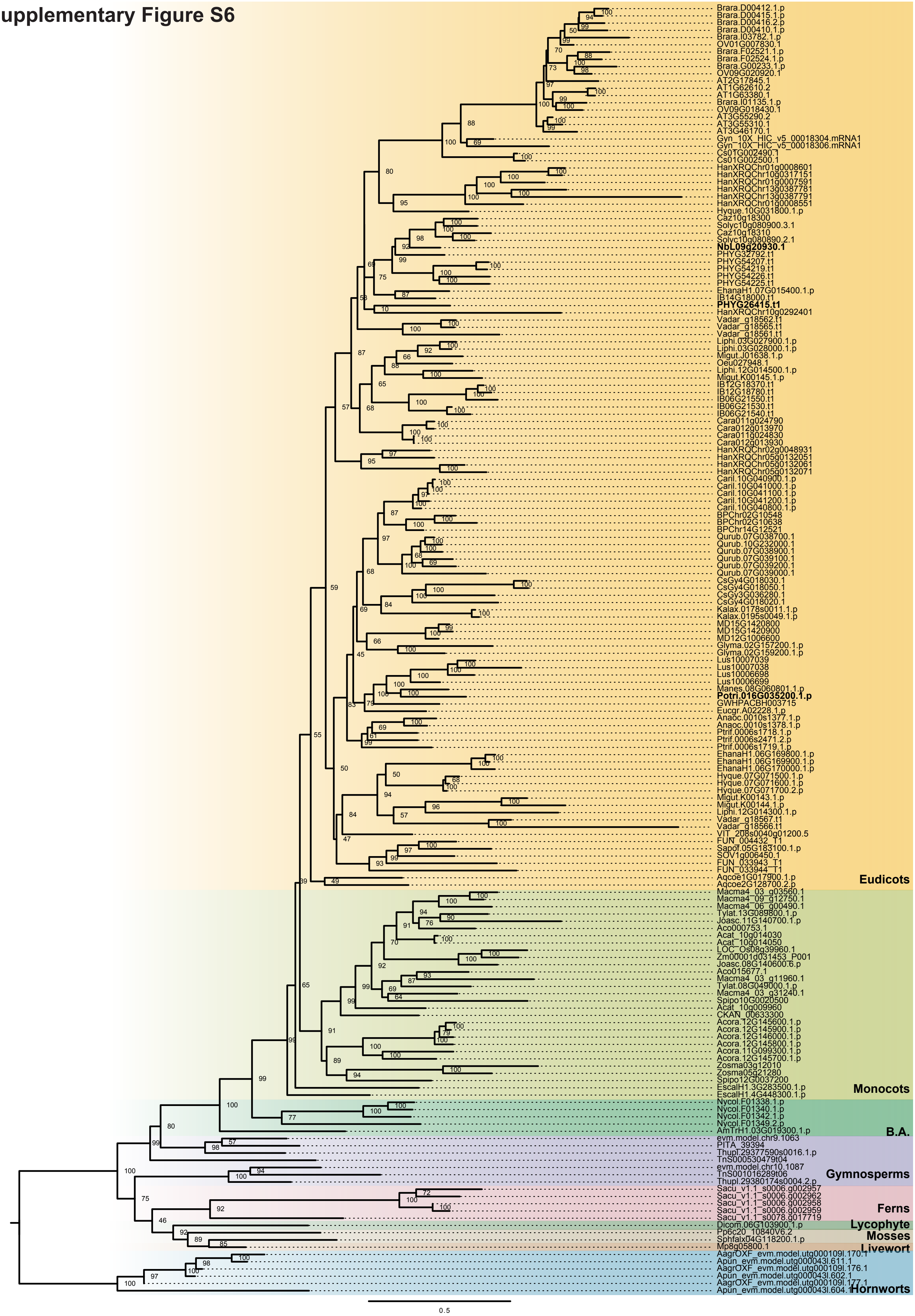


**Supplementary Figure S6. Phylogeny of *Nb*BalSα and its homologs in plants.**
Phylogenetic tree of BalSα homologs was constructed using protein sequences retrieved from the BLASTp searches across 84 representative plant species spanning major Archaeplastida lineages. Sequences were aligned using MAFFT (v7.490), and the tree was built using IQ-TREE (v2.2.2.6) under the maximum-likelihood method with 1,000 ultrafast bootstrap replicates. The resulting tree was visualized and annotated in FigTree (v1.4.4). Ultrafast bootstrap support values (%) are shown next to nodes. Reported genes encoding BalSα are highlighted in bold. B.A., basal angiosperms.

### Supplementary Figure S7

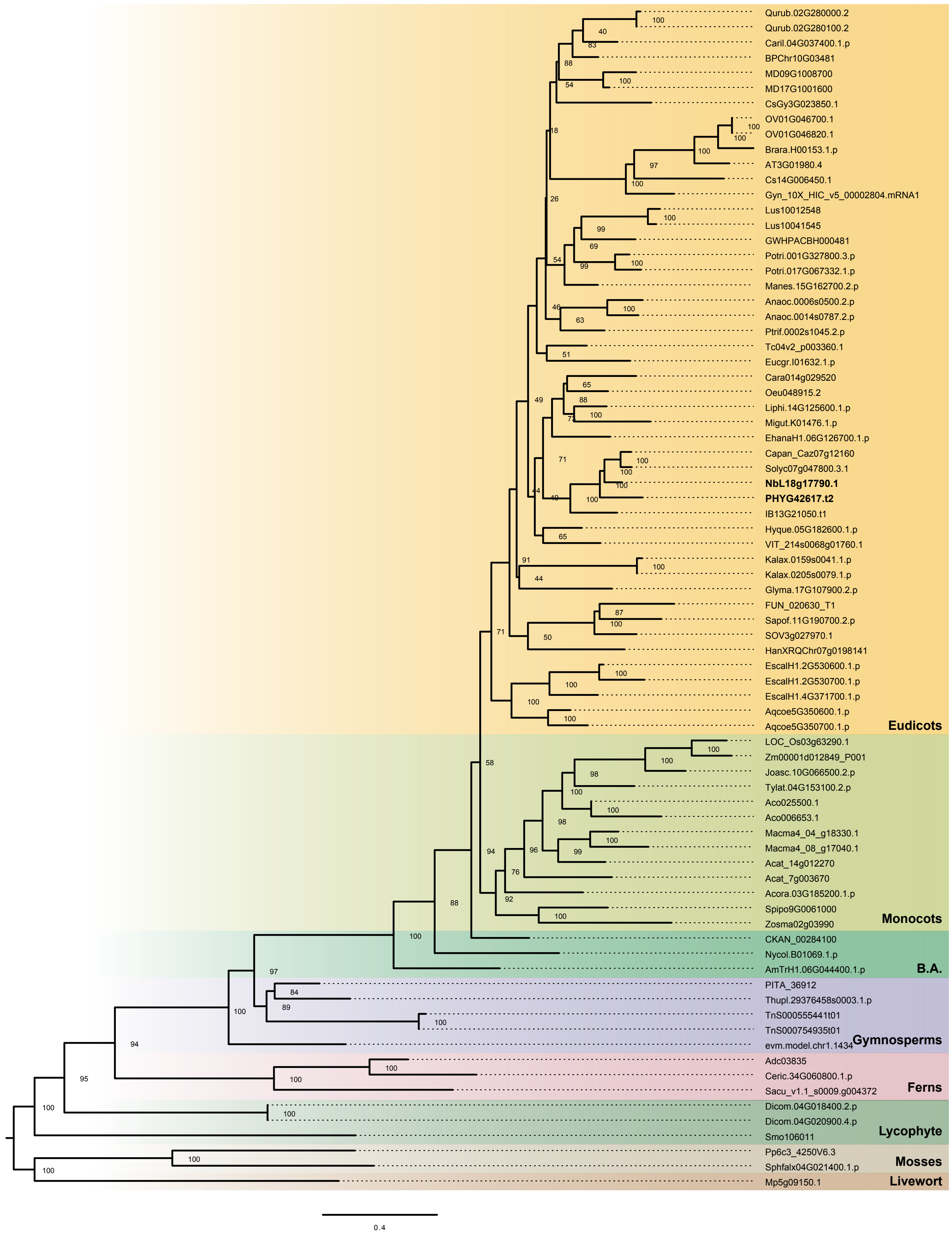


**Supplementary Figure S7. Phylogeny of *Nb*BalSβ and its homologs in plants.**
Phylogenetic tree of BalSβ homologs was constructed using protein sequences retrieved from the BLASTp searches across 84 representative plant species spanning major Archaeplastida lineages. Sequences were aligned using MAFFT (v7.490), and the tree was built using IQ-TREE (v2.2.2.6) under the maximum-likelihood method with 1,000 ultrafast bootstrap replicates. The resulting tree was visualized and annotated in FigTree (v1.4.4). Ultrafast bootstrap support values (%) are shown next to nodes. Reported gene encoding BalSβ is highlighted in bold. B.A., basal angiosperms.

#### Supplementary Figure S8

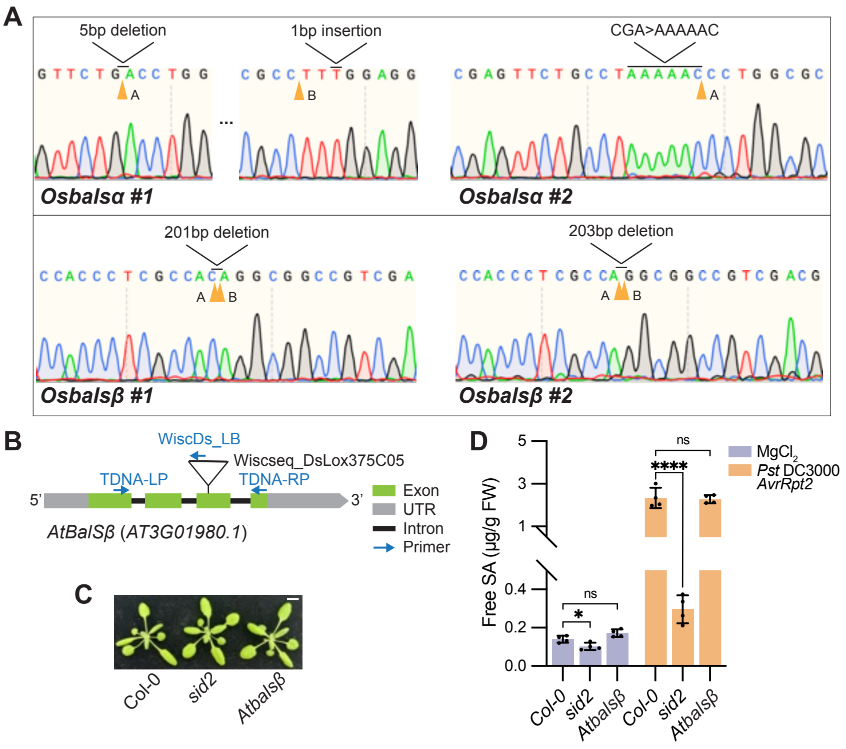


**Supplementary Figure S8**. **Validation of rice CRISPR lines by Sanger sequencing and analysis of *Arabidopsis balsβ* mutants.**

**A)** Sanger sequencing chromatograms of two independent mutants of each gene, *OsBalSα* and *OsBalSβ*. Line *Osbalsα #1* carries a 5-bp deletion at sgRNA target site A and a 1-bp insertion at site B. *Osbalsα #2* has a substitution/insertion replacing CGA with AAAAAC at site A. *Osbalsβ #1* and *Osbalsβ #2* harbor 201-bp and 203-bp deletions between sites A and B, respectively.

**B)** Structure of the single *AtBalSβ* gene showing location of the T-DNA insertion in the mutant line *Atbalsβ*. Exons are represented by green rectangles, UTRs by grey rectangles, and introns by connecting black lines. The primers used for PCR genotyping are depicted with blue arrows.

**C)** Morphology of three-week-old Col-0, *sid2* (mutating *ICS1*) and *Atbalsβ* plants grown under long-day conditions. Scale bar, 1 cm.

**D)** Free SA levels of four-week-old soil-grown *Arabidopsis* plants of the indicated genotypes at 16 hpi with *Pst* DC3000 avrRpt2 ($OD_{600}$ = 0.005) or $MgCl_2$, as measured by HPLC. Asterisks indicate statistically significant differences (two-tailed Student's *t*-tests, **P* < 0.05, n = 4 independent plants). ns, no significant difference.

### Supplementary Figure S9

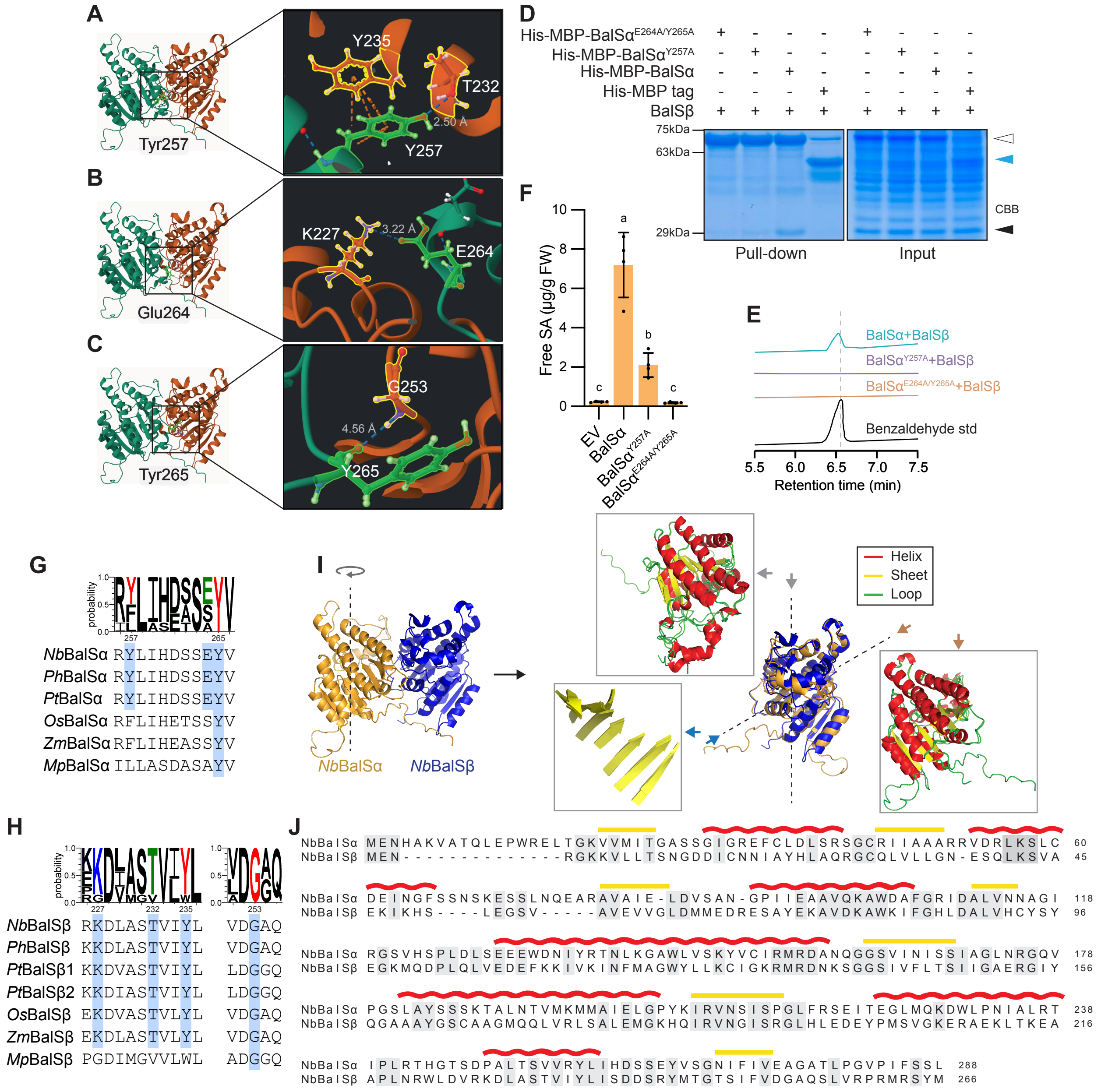


**Supplementary Figure S9. Structural analysis of the BalSα-BalSβ interface.**
**A-C)** Ribbon diagrams of the BalSα-BalSβ heterodimer highlighting predicted interface residues. Tyr257 in BalSα was predicted to simultaneously form a hydrogen bond with Thr232 and engage in π-π stacking with Tyr235 of BalSβ, whereas Glu264 and Tyr265 were predicted to form inter-chain hydrogen bonds with Lys227 and Gly253 of BalSβ, respectively. Zoom-in views show BalSα residues Tyr257 (**A**), Glu264 (**B**), and Tyr265 (**C**) (green sticks) and their interacting residues in BalSβ (orange sticks), with distances indicated. Hydrogen bonds (blue dashed lines) and π-π stacking interactions (orange dashed lines) were identified using RING.
**D)** Coomassie-stained SDS-PAGE gel from *in vitro* pull-down assays showing co-purification of untagged BalSβ with 6×His-MBP-tagged BalSα, its mutant variants or the tag alone using Ni-NTA resin. Proteins were expressed in *E. coli*, equal amounts of lysates were incubated with the same amount of BalSβ and Ni-NTA beads, and eluates were analyzed by SDS-PAGE. Filled triangle indicates BalSβ (~29 kDa), open triangle indicates His-MBP-BalSα and its variants (~74 kDa), and blue triangle indicates the His-MBP control (~43 kDa).
**E)** *In vitro* benzoyl-CoA reduction into benzaldehyde by BalSβ together with BalSα, BalSα$^{Y257A}$ or BalSα$^{E264A/Y265A}$. Reactions containing BalSβ and BalSα or its variants were incubated with benzyl-CoA and NADPH at 28 °C for 30 min. Benzaldehyde formation was verified by HPLC-DAD against an authentic standard.
F) Free SA levels in balsα mutants transiently overexpressed with empty vector (EV), BalSα, BalSα$^{Y257A}$ or BalSα$^{E264A/Y265A}$ at 8 hpi with *Pst* DC3000 ($OD_{600}$ = 0.02), as measured by a biosensor-based method. *N. benthamiana* leaves were infiltrated with *Agrobacterium* carrying the overexpression constructs ($OD_{600}$ = 0.4) three days before *Pst* DC3000 inoculation. Error bars represent mean ± s.d. (one-way ANOVA with Tukey's test, n = 4 independent plants). Letters indicate statistically significant differences.
**G-H)** Sequence alignment of regions surrounding key interface residues in BalSα (**G**) and BalSβ (**H**) across representative species. Identical residues are highlighted in blue.
**I)** Ribbon diagram of BalS heterodimer and superimposition of the two subunits showing high structural symmetry. BalSα (yellow) and BalSβ (blue) exhibit strong structural overlap (RMSD = 0.737 Å). Secondary structural elements are colored as follows: α-helices (red), β-sheets (yellow), and loops (green). Insets show top, front, and back views of the superimposed subunits. Models were generated with AlphaFold2 and visualized in PyMOL.
**J)** Sequence alignment of BalSα and BalSβ. Predicted secondary structure elements based on AlphaFold are shown above the alignment (α-helices in red and β-sheets in yellow). Conserved residues are indicated by grey shading.

### Supplementary Figure S10

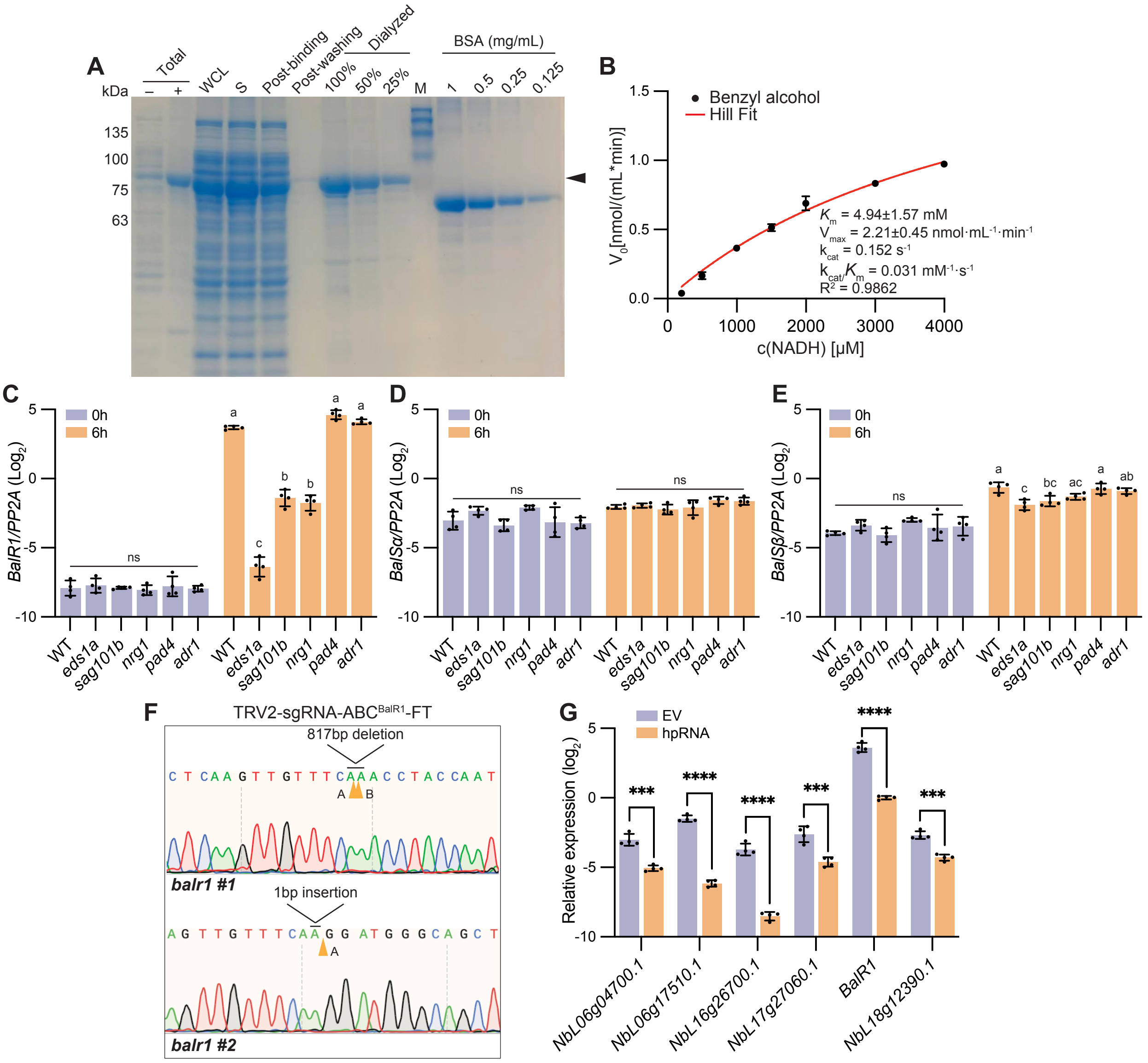


**Supplementary Figure S10. Functional characterization and regulation of *Nb*BalR1 in SA biosynthesis.**

**A)** Coomassie-stained SDS-PAGE gel showing purification of His-MBP-tagged BalR1 expressed in *E. coli*. Total, bacterial cultures with (+) or without (-) IPTG induction; WCL, whole cell lysate; S, soluble lysate after sonication; Post-binding, flow-through after Ni-NTA binding; Post-washing, flow-through after washing; Dialyzed: fractions after dialysis. M, marker. Black triangle indicates purified His-MBP tagged BalR1 (~82 kDa). BSA standards were used to estimate protein concentration.

**B)** Kinetic analysis of BalR1 activity with varying concentrations of NADH in the presence of benzaldehyde. Data from three replicates were fitted using non-linear Hill regression in Prism 10 (v10.2.0).

**C-E, G)** Relative expression of *NbBalR1* (**C**), *NbBalSα* (**D**), and *NbBalSβ* (**E**) in the indicated genotypes at 6 hpi with *Pst* DC3000 ($OD_{600}$ = 0.02), and of *ADH* genes in hpRNA-silenced plants relative to empty vector (EV) controls (**G**), measured by qRT–PCR ($log_2$ scale). Error bars represent mean ± s.d. ns, no significant difference. Letters indicate statistically significant differences (one-way ANOVA with Tukey's test, n = 4 independent plants). Asterisks indicate statistically significant differences (two-tailed Student's *t*-tests, *$P < 0.05$, n = 4 independent plants).

**F)** Sanger sequencing chromatograms of two independent CRISPR mutants of *NbBalR1*. Line *balr1 #1* carries an 817-bp deletion between sgRNA target site A and B. Line *balr1 #2* carries a 1-bp insertion at site A.

### Supplementary Figure S11

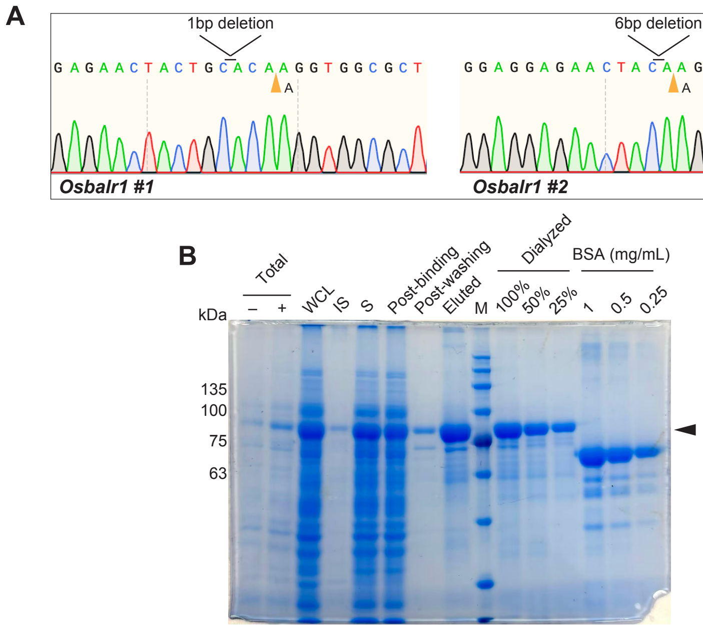


**Supplementary Figure S11**. **Validation of rice CRISPR lines by Sanger sequencing and purification of *Os*BalR1.**

**A)** Sanger sequencing chromatograms of two independent mutants of *OsBalR1*. Line *Osbalr1 #1* and line *Osbalr1 #2* carry 1-bp and 6-bp deletions at target site A, respectively.

**B)** Coomassie-stained SDS-PAGE gel showing the purified *Os*BalR1 expressed in *E. coli*. Total, bacterial cultures with (+) or without (-) IPTG induction; WCL, whole cell lysate; IS, insoluble fraction; S, soluble lysate after sonication; Post-binding, flow-through after Ni-NTA binding; Post-washing, flow-through after washing; Eluted, eluate; Dialyzed: fractions after dialysis. M, marker. Black triangle indicates purified His-MBP tagged *Os*BalR1 (~82 kDa). BSA standards were used to estimate protein concentration.
