## Supplemental Tables for "Identification of Reductases Catalyzing Benzyl Alcohol Formation during Salicylic Acid Biosynthesis in Plants"

### Supplementary Table 1. Candidate genes identified by whole-genome sequencing (WGS).

| Position (bp) | Gene | Gene/Region | Allele count (Mut/WT) | Allele frequency |
|---|---|---|---|---|
| **4-66** | | | | |
| 107725048 | *NbL09g17410.1* | Exonic (nonsynonymous) | 21/3 | 0.87 |
| 109420660 | *NbL09g17660.1* | Intronic | 15/2 | 0.88 |
| 114462776 | *NbL09g18460.1–NbL09g18470.1* | Intergenic | 3/0 | 1.00 |
| 124320737 | *NbL09g19940.1–NbL09g19950.1* | Intergenic | 23/0 | 1.00 |
| 124737678 | *NbL09g20020.1–NbL09g20030.1* | Intergenic | 24/0 | 1.00 |
| 126783070 | *NbL09g20390.1–NbL09g20400.1* | Intergenic | 25/0 | 1.00 |
| 129042432 | *NbL09g20770.1* | Downstream | 21/0 | 1.00 |
| 130053953 | *NbL09g20930.1* | Exonic (nonsynonymous) | 14/0 | 1.00 |
| 131332856 | *NbL09g21170.1–NbL09g21180.1* | Intergenic | 25/1 | 0.96 |
| 138597817 | *NbL09g22260.1* | Downstream | 24/2 | 0.92 |
| 138624536 | *NbL09g22280.1* | Intronic | 28/5 | 0.85 |
| 141790164 | *NbL09g23040.1–NbL09g23050.1* | Intergenic | 12/2 | 0.86 |
| **15-328** | | | | |
| 107724172 | *NbL09g17410.1* | Exonic (nonsynonymous) | 22/4 | 0.85 |
| 110576743 | *NbL09g17820.1* | Upstream | 18/4 | 0.82 |
| 112721375 | *NbL09g18200.1* | Exonic (nonsynonymous) | 13/3 | 0.81 |
| 118955758 | *NbL09g19080.1* | Downstream | 12/0 | 1.00 |
| 119108368 | *NbL09g19100.1* | Upstream | 18/0 | 1.00 |
| 120516520 | *NbL09g19330.1* | UTR3 | 17/0 | 1.00 |
| 121206893 | *NbL09g19480.1* | Downstream | 16/1 | 0.94 |
| 124415846 | *NbL09g19960.1* | Downstream | 22/1 | 0.96 |
| 125993925 | *NbL09g20300.1* | Downstream | 9/0 | 1.00 |
| 129170419 | *NbL09g20780.1* | Exonic (nonsynonymous) | 24/0 | 1.00 |
| 130051521 | *NbL09g20930.1* | Exonic (nonsynonymous) | 17/0 | 1.00 |
| 131369924 | *NbL09g21190.1* | Exonic (unknown) | 23/0 | 1.00 |
| 133965652 | *NbL09g21580.1* | Upstream | 11/0 | 1.00 |
| 134762681 | *NbL09g21730.1* | Upstream | 23/0 | 1.00 |
| 138488324 | *NbL09g22220.1* | UTR5 | 25/0 | 1.00 |
| 138622999 | *NbL09g22280.1* | Downstream | 20/1 | 0.95 |
| 139985032 | *NbL09g22630.1* | Downstream | 14/1 | 0.93 |
| 140550396 | *NbL09g22770.1* | UTR3 | 20/1 | 0.95 |
| 142749279 | *NbL09g23290.1* | UTR3 | 15/3 | 0.83 |
| **13-270** | | | | |
| 118170755 | *NbL18g17130.1* | UTR3 | 28/1 | 0.97 |
| 118384875 | *NbL18g17180.1* | Downstream | 23/0 | 1.00 |
| 118494084 | *NbL18g17190.1* | Intronic | 15/0 | 1.00 |
| 118682925 | *NbL18g17220.1* | UTR3 | 26/1 | 0.96 |
| 119713663 | *NbL18g17350.1* | Intergenic | 22/1 | 0.96 |
| 120452743 | *NbL18g17490.1* | Upstream | 29/0 | 1.00 |
| 120669348 | *NbL18g17530.1* | Intronic | 19/0 | 1.00 |
| 122253471 | *NbL18g17790.1* | Splicing | 24/0 | 1.00 |
| 122323002 | *NbL18g17810.1* | Intronic | 4/0 | 1.00 |
| 124946415 | *NbL18g17850.1* | Intronic | 14/1 | 0.93 |
| 123852741 | *NbL18g18070.1* | Upstream | 21/1 | 0.95 |
| 124549231 | *NbL18g18190.1* | Intronic | 4/1 | 0.80 |
| 125823905 | *NbL18g18350.1* | Exonic (nonsynonymous) | 21/0 | 1.00 |
| 126585176 | *NbL18g18420.1* | Downstream | 19/0 | 1.00 |
| 127884513 | *NbL18g18610.1* | Intronic | 25/1 | 0.96 |
| 128111435 | *NbL18g18660.1* | Downstream | 13/2 | 0.87 |
| 130215835 | *NbL18g18910.1* | Exonic (nonsynonymous) | 22/2 | 0.92 |

### Supplementary Table 2. Sources and database versions used for phylogenetic analysis.

| Classes | Species | BalSα | BalSβ | Source | Version |
|---|---|---|---|---|---|
| Rhodophyta | *Cyanidioschyzon merolae* | 0 | 0 | JGI | Soos |
| | *Galdieria sulphuraria* | 0 | 0 | JGI | Azora |
| | *Porphyra umbilicalis* | 0 | 0 | JGI | 4086291 |
| | *Chondrus crispus* | 0 | 0 | JGI | Stackhouse |
| Chlorophyta | *Bathycoccus prasinos* | 0 | 0 | JGI | RCC1105 |
| | *Chlamydomonas eustigma* | 0 | 0 | JGI | NIES-2499 |
| | *Chlamydomonas reinhardtii* | 0 | 0 | JGI | CC-4532 v6.1 |
| | *Chlorella variabilis* | 0 | 0 | JGI | NC64A v1.0 |
| | *Coccomyxa subellipsoidea* | 0 | 0 | JGI | C-169 v3.0 |
| | *Gonium pectorale* | 0 | 0 | JGI | NIES-2863 |
| | *Micromonas commoda* | 0 | 0 | JGI | NOUM17 (RCC 299) |
| | *Micromonas pusilla* | 0 | 0 | JGI | CCMP1545 |
| | *Monoraphidium neglectum* | 0 | 0 | JGI | SAG 48.87 |
| | *Volvox carteri* | 0 | 0 | JGI | v2.1 |
| Streptophyte algae | *Mesostigma viride* | 0 | 0 | JGI | NIES-296 |
| | *Klebsormidium nitens* | 0 | 0 | JGI | v1.1 |
| | *Chara braunii* | 0 | 0 | JGI | S276 |
| Hornworts | *Anthoceros agrestis* | 3 | 0 | UZH | Oxford |
| | *Anthoceros punctatus* | 3 | 0 | UZH | Anthoceros punctatus PROT |
| Liverworts | *Marchantia polymorpha* | 1 | 1 | MarpoBase | MpTak-1 v7.1 |
| Mosses | *Physcomitrella patens* | 1 | 1 | PhytozomeV14 | v6.1 |
| | *Sphagnum fallax* | 1 | 1 | PhytozomeV14 | v1.1 |
| Lycophytes | *Selaginella moellendorffii* | 0 | 1 | PhytozomeV14 | v1.0 |
| | *Diphasiastrum complanatum* | 1 | 2 | PhytozomeV14 | v3.1 |
| Ferns | *Salvinia cucullata* | 5 | 1 | FernBase | v1.2 |
| | *Ceratopteris richardii* | 0 | 1 | NCBI | v2 |
| | *Adiantum capillus-veneris* | 0 | 1 | NCBI | ASM1452938v2 |
| Gymnosperms | *Gnetum montanum* | 2 | 2 | TreeGenes | v1.0 |
| | *Thuja plicata* | 2 | 1 | PhytozomeV14 | v3.1 |
| | *Pinus taeda* | 1 | 1 | TreeGenes | v2.01 |
| | *Ginkgo biloba* | 2 | 1 | GinkgoDB | Ver. 2021 |
| Basal Angiosperm | *Amborella trichopoda* | 1 | 1 | PhytozomeV14 | v2.1 |
| | *Nymphaea colorata* | 4 | 1 | PhytozomeV14 | v1.2 |
| | *Cinnamomum kanehirae* | 1 | 1 | PhytozomeV14 | v3 |
| Monocots | *Zostera marina* | 2 | 1 | PhytozomeV14 | v3.1 |
| | *Spirodela polyrhiza* | 2 | 1 | PhytozomeV14 | v2 |
| | *Acorus americanus* | 6 | 1 | PhytozomeV14 | v1.1 |
| | *Spirodela polyrhiza* | 2 | 1 | PhytozomeV14 | v2 |
| | *Zostera marina* | 2 | 1 | PhytozomeV14 | v3.1 |
| | *Areca catechu* | 3 | 2 | NCBI | v1 |
| | *Typha latifolia* | 2 | 1 | PhytozomeV14 | v1.1 |
| | *Joinvillea ascendens* | 2 | 1 | PhytozomeV14 | v1.1 |
| | *Zea mays* | 1 | 1 | PhytozomeV14 | v4 |
| | *Oryza sativa* | 1 | 1 | PhytozomeV14 | v7.0 |
| | *Musa acuminata* | 5 | 2 | Banana Genome Hub | v4 |
| | *Ananas comosus* | 3 | 2 | PhytozomeV14 | v3 |
| Eudicots | *Eschscholzia californica* | 2 | 3 | NCBI | dmEscCali1.0.p |
| | *Aquilegia coerulea* | 2 | 2 | PhytozomeV14 | v3.1 |
| | *Kalanchoe laxiflora* | 2 | 2 | PhytozomeV14 | v1.1 |
| | *Vitis vinifera* | 1 | 1 | PhytozomeV14 | v2.1 |
| | *Populus trichocarpa* | 1 | 2 | PhytozomeV14 | v4.1 |
| | *Linum usitatissimum* | 4 | 2 | PhytozomeV14 | v1 |
| | *Kandelia obovata* | 1 | 1 | NCBI | v1 |
| | *Manihot esculenta* | 1 | 1 | NCBI | v8.1 |
| | *Glycine max* | 2 | 1 | PhytozomeV14 | Wm82.a6.v1 |
| | *Malus domestica* | 3 | 2 | NCBI | GDT2T_hap1 |
| | *Cucumis sativus* | 4 | 1 | NCBI | 9930_V3 |
| | *Quercus rubra* | 6 | 2 | PhytozomeV14 | v2.1 |
| | *Betula platyphylla* | 3 | 1 | PhytozomeV14 | v1.1 |
| | *Carya illinoinensis* | 5 | 1 | PhytozomeV14 | v1.1 |
| | *Eucalyptus grandis* | 1 | 1 | PhytozomeV14 | v2.0 |
| | *Poncirus trifoliata* | 3 | 1 | PhytozomeV14 | v1.3.1 |
| | *Anacardium occidentale* | 2 | 2 | PhytozomeV14 | v0.9 |

| | | | | | |
|---|---|---|---|---|---|
| | *Capparis spinosa* | 2 | 1 | (Wang et al., 2022) | - |
| | *Gynandropsis gynandra* | 2 | 1 | (Hoang et al., 2023) | v3.0 |
| | *Orychophragmus violaceus* | 3 | 2 | (Zhang et al., 2023) | - |
| | *Arabidopsis thaliana* | 6 | 1 | PhytozomeV14 | Araport11 |
| | *Brassica rapa* | 9 | 1 | PhytozomeV14 | v1.3 |
| | *Spinacia oleracea* | 1 | 1 | PhytozomeV14 | Spov3 |
| | *Saponaria officinalis* | 1 | 1 | PhytozomeV14 | v1.1 |
| | *Portulaca amilis* | 3 | 1 | PhytozomeV14 | v1.0 |
| | *Hydrangea quercifolia* | 4 | 1 | PhytozomeV14 | v1.1 |
| | *Vaccinium darrowii* | 5 | 1 | PhytozomeV14 | v1.2 |
| | *Ipomoea batatas* | 6 | 1 | Ipomoea Genome Hub | v1.0.a2 |
| | *Capsicum annuum* | 2 | 1 | PepperGD | Zhangshugang |
| | *Petunia axillaris* | 6 | 1 | NCBI | ASM2999057v1 |
| | *Nicotiana benthamiana* | 1 | 1 | apollo.nbenth | LAB360 |
| | *Solanum lycopersicum* | 2 | 1 | PhytozomeV14 | ITAG5.0 |
| | *Coffea arabica* | 4 | 1 | (Salojärvi et al., 2024) | - |
| | *Mimulus guttatus* | 4 | 1 | PhytozomeV14 | v2.0 |
| | *Olea europaea* | 1 | 1 | PhytozomeV14 | v1.0 |
| | *Lindenbergia philippensis* | 4 | 1 | PhytozomeV14 | v1.1 |
| | *Ehretia anacua* | 4 | 1 | JGI | HAP1 v1.1 |
| | *Helianthus annuus* | 11 | 1 | PhytozomeV14 | r1.2 |

### Supplementary Table 3. Expression of annotated dehydrogenases among XopQ-responsive transcripts.

| | |
|---|---|
| logFC | Log2 scale of fold changes of the gene between the comparison groups |
| logCPM | Log2 scale of the average counts per milion of the gene in the comparison groups |
| F | Quasi-likelihood F-test score |
| FDR | False discovery rate |

| Transcript ID | Annotation | Nb-XopQ/Nb-EV | | | | |
|---|---|---|---|---|---|---|
| | | **logFC** | **logCPM** | **F** | **PValue** | **FDR** |
| Nbv5.1tr6236361 | Proline dehydrogenase 2, mitochondrial (probable) | 6.11452492 | 1.51645506 | 97.6127322 | 1.08E-15 | 3.41E-13 |
| **Nbv5.1tr6207848** | **Probable cinnamyl alcohol dehydrogenase 6 (probable)** | **4.40937338** | **4.01399815** | **115.876116** | **1.88E-17** | **9.58E-15** |
| Nbv5.1tr6203320 | Arogenate dehydrogenase 1, chloroplastic (probable) | 3.88903551 | -0.3442597 | 19.8422334 | 2.59E-05 | 0.0003915 |
| Nbv5.1tr6223703 | L-lactate dehydrogenase A (probable) | 3.54521409 | 0.11659234 | 25.2268969 | 2.85E-06 | 5.52E-05 |
| Nbv5.1tr6203866 | Dehydrogenase/reductase SDR family member 4 (probable) | 2.90615868 | 2.70792803 | 59.7083645 | 2.20E-11 | 2.04E-09 |
| Nbv5.1tr6229542 | Dihydrolipoyl dehydrogenase 1, mitochondrial (probable) | 2.73284061 | -0.2289776 | 14.0075821 | 0.00033337 | 0.00349151 |
| Nbv5.1tr6211998 | 6-phosphogluconate dehydrogenase, decarboxylating 2 (probable) | 2.68240052 | -0.5183667 | 11.1124218 | 0.00128018 | 0.01063755 |
| Nbv5.1tr6219156 | Dihydrolipoyllysine-residue succinyltransferase component of 2-oxoglutarate dehydrogenase complex 1, mitochondrial (probable) | 2.66620725 | 0.52347854 | 24.0558801 | 4.56E-06 | 8.32E-05 |
| Nbv5.1tr6206328 | 6-phosphogluconate dehydrogenase, decarboxylating (probable) | 2.63359421 | 1.68893529 | 40.9508458 | 8.63E-09 | 3.24E-07 |
| Nbv5.1tr6225092 | 6-phosphogluconate dehydrogenase, decarboxylating (probable) | 2.60890373 | -0.8449626 | 7.99166531 | 0.00588002 | 0.03532636 |
| Nbv5.1tr6207765 | (+)-neomenthol dehydrogenase (probable) | 2.53962707 | 1.22793137 | 32.8591114 | 1.53E-07 | 4.25E-06 |
| Nbv5.1tr6244019 | Isovaleryl-CoA dehydrogenase 2, mitochondrial (probable) | 2.37763018 | 0.05193765 | 13.9521501 | 0.00034188 | 0.00356297 |
| Nbv5.1tr6236624 | Isocitrate dehydrogenase [NADP] (probable) | 2.04891788 | 0.74416724 | 16.5377858 | 0.00010754 | 0.00132016 |
| Nbv5.1tr6212047 | Dihydrolipoyllysine-residue succinyltransferase component of 2-oxoglutarate dehydrogenase complex 1, mitochondrial (probable) | 1.92944902 | 0.10864941 | 10.3478499 | 0.00184533 | 0.01425551 |
| Nbv5.1tr6216442 | Arogenate dehydrogenase 2, chloroplastic (probable) | 1.85827215 | 1.67002244 | 24.6765862 | 3.55E-06 | 6.68E-05 |
| Nbv5.1tr6205871 | Dihydrolipoyllysine-residue succinyltransferase component of 2-oxoglutarate dehydrogenase complex 1, mitochondrial (probable) | 1.70543594 | 2.92671719 | 24.731282 | 3.47E-06 | 6.56E-05 |
| Nbv5.1tr6243652 | Isocitrate dehydrogenase [NADP] (similar to) | 1.58994302 | 0.06838984 | 7.4856582 | 0.00759592 | 0.04286248 |
| Nbv5.1tr6237566 | Probable mannitol dehydrogenase (probable) | 1.58438063 | 0.31868246 | 7.79756488 | 0.00648472 | 0.03808808 |
| Nbv5.1tr6203974 | Succinate dehydrogenase [ubiquinone] iron-sulfur subunit 2, mitochondrial (probable) | 1.35042081 | 2.49885669 | 15.5252697 | 0.00016834 | 0.00193958 |
| Nbv5.1tr6221174 | Glutamate dehydrogenase B (probable) | 1.33154113 | 1.60201283 | 12.5683947 | 0.00064597 | 0.00602934 |
| Nbv5.1tr6219074 | 2-oxoglutarate dehydrogenase, mitochondrial (probable) | 1.29511383 | 0.52714168 | 7.149178 | 0.00902027 | 0.04902386 |
| Nbv5.1tr6225820 | Pyruvate dehydrogenase E1 component subunit alpha, mitochondrial (probable) | 1.13670009 | 1.41487571 | 8.89844959 | 0.00374117 | 0.0248475 |
| Nbv5.1tr6220224 | Isocitrate dehydrogenase [NADP], chloroplastic (probable) | -1.312609 | 0.48749341 | 7.62470164 | 0.00707785 | 0.04077888 |
| Nbv5.1tr6229700 | Betaine aldehyde dehydrogenase, chloroplastic (probable) | -1.3344921 | 1.65576912 | 14.2271005 | 0.00030173 | 0.00322037 |
| Nbv5.1tr6205758 | Aldehyde dehydrogenase family 2 member B4, mitochondrial (probable) | -1.3799521 | 2.53168314 | 16.6078453 | 0.00010428 | 0.00128416 |
| Nbv5.1tr6204482 | Probable NADH dehydrogenase [ubiquinone] 1 alpha subcomplex subunit 5, mitochondrial (probable) | -1.4017003 | 0.49851859 | 8.64863953 | 0.004234 | 0.02745727 |

| | | | | | | |
|---|---|---|---|---|---|---|
| Nbv5.1tr6243702 | NADP-dependent glyceraldehyde-3-phosphate dehydrogenase (probable) | -1.409067 | 2.17142036 | 16.4778321 | 0.00011041 | 0.00134884 |
| Nbv5.1tr6229525 | Glyceraldehyde-3-phosphate dehydrogenase B, chloroplastic (probable) | -1.6532893 | 2.15177885 | 17.7751365 | 6.27E-05 | 0.00084161 |
| Nbv5.1tr6233580 | Malate dehydrogenase [NADP], chloroplastic (probable) | -1.7571723 | 0.4811897 | 11.3276668 | 0.00115591 | 0.00976423 |
| Nbv5.1tr6230639 | L-idonate 5-dehydrogenase (probable) | -2.136735 | 0.35376653 | 16.5008049 | 0.0001093 | 0.00133819 |
| Nbv5.1tr6205686 | Glyceraldehyde-3-phosphate dehydrogenase B, chloroplastic (probable) | -2.212115 | -0.3989603 | 9.9916097 | 0.00219167 | 0.01639211 |
| Nbv5.1tr6219998 | Glucose-6-phosphate 1-dehydrogenase, chloroplastic (probable) | -2.4096946 | 0.61117436 | 24.405008 | 3.96E-06 | 7.35E-05 |
| Nbv5.1tr6204880 | Glyceraldehyde-3-phosphate dehydrogenase A, chloroplastic (probable) | -2.4674072 | 5.84625678 | 50.0855014 | 4.22E-10 | 2.42E-08 |
| Nbv5.1tr6205685 | Glyceraldehyde-3-phosphate dehydrogenase B, chloroplastic (probable) | -2.5383873 | 0.29195332 | 19.1496453 | 3.47E-05 | 0.00050956 |
| Nbv5.1tr6214416 | Glycerol-3-phosphate dehydrogenase [NAD(P)+] (probable) | -2.85042 | -0.3585749 | 16.8219109 | 9.49E-05 | 0.00119263 |
| Nbv5.1tr6230169 | Glyceraldehyde-3-phosphate dehydrogenase B, chloroplastic (probable) | -3.0299241 | 2.11066952 | 56.7340603 | 5.36E-11 | 4.27E-09 |
| Nbv5.1tr6225311 | Malate dehydrogenase [NADP], chloroplastic (probable) | -3.1287116 | -0.4983874 | 16.8982518 | 9.18E-05 | 0.00116386 |
| Nbv5.1tr6229548 | Glycerol-3-phosphate dehydrogenase [NAD(+)] (probable) | -3.6473985 | -0.8237365 | 16.3223613 | 0.00011824 | 0.00142887 |

### Supplementary Table 4. The top 50 co-expressed genes of *OsBalR1* (*Os04g15920.1*).

| Known pathway gene | Rank | Gene | Function | Other ID | osa-u.5 for 4335223 | osa-r.6 for 4335223 | osa-m.8 for 4335223 | osa-e.1 for 4335223 |
|---|---|---|---|---|---|---|---|---|
| *OsCNL* | 1 | LOC4331500 | trans-cinnamate:CoA ligase, peroxisomal | Os03g0130100 | 7.1 | 5.9 | 8.3 | 10.4 |
| | 2 | LOC4347069 | WRKY transcription factor WRKY76-like | Os09g0417600 | 5.2 | 4.9 | 5.6 | 5.5 |
| | 3 | LOC4324018 | NRR repressor homolog 2-like | Os01g0508500 | | | | |
| *OsKAT2* | 4 | LOC4348804 | 3-ketoacyl-CoA thiolase 2, peroxisomal | Os10g0457600 | 5 | 4.6 | 5.4 | 3 |
| *OsBBO* | 5 | LOC4347176 | trimethyltridecatetraene synthase | Os09g0441400 | 5 | 5.8 | 4 | 2.6 |
| | 6 | LOC4347070 | WRKY transcription factor WRKY62-like | Os09g0417800 | 4.7 | 4 | 5.3 | 3.4 |
| | 7 | LOC4328124 | S-(+)-linalool synthase, chloroplastic-like | OSNPB_020121700 | 4.7 | 3.2 | 6.1 | 2.6 |
| *OsBEBT* | 8 | LOC4349050 | benzyl alcohol O-benzoyltransferase | Os10g0503300 | 4.6 | 6.8 | 1.4 | 6.6 |
| | 9 | LOC9272383 | geranylgeranyl pyrophosphate synthase 7, chloroplastic | Os01g0248701 | 4.5 | 6.7 | | 4.6 |
| *OsPAL7* | 10 | LOC4336415 | phenylalanine ammonia-lyase | Os04g0518400 | 4.5 | 5.2 | 3.8 | 2.5 |
| | 11 | LOC4338413 | transcription factor WRKY45-1-like | Os05g0322900 | 4.2 | 3.9 | 4.5 | 3.3 |
| | 12 | LOC4330542 | anthranilate O-methyltransferase 1 | Os02g0719600 | 4.1 | 2.7 | 5.5 | 5 |
| | 13 | LOC4335756 | arogenate dehydratase/prephenate dehydratase 6, chloroplastic | Os04g0406600 | 4.1 | 3.6 | 4.5 | 3.1 |
| | 14 | LOC4330040 | phenylalanine ammonia-lyase-like | OSNPB_020627100 | 4.5 | 5.2 | 3.8 | 2.5 |
| | 15 | LOC4336765 | protein DMR6-LIKE OXYGENASE 1 | Os04g0581100 | 3.7 | 3 | 4.4 | 2 |
| | 16 | LOC4345658 | probable glucomannan 4-beta-mannosyltransferase 11 | Os08g0434632 | 3.7 | 2.9 | 4.6 | 3.1 |
| | 17 | LOC4342614 | probable 1-deoxy-D-xylulose-5-phosphate synthase 2, chloroplastic | OSNPB_070190000 | 3.7 | 4.4 | 3 | 2.3 |
| | 18 | LOC4338644 | annexin D4 | Os05g0382600 | 3.6 | 3.8 | 3.4 | 1.9 |
| | 19 | LOC4343852 | protein EGG APPARATUS-1 | Os07g0605400 | 3.5 | 3.3 | 3.8 | 2.5 |
| | 20 | LOC4343946 | phospho-2-dehydro-3-deoxyheptonate aldolase 2, chloroplastic-like | OSNPB_070622200 | 3.5 | 3.7 | 3.3 | 0.8 |
| | 21 | LOC4334549 | calmodulin-like protein 2 | Os03g0812400 | 3.4 | 3.4 | 3.3 | |
| | 22 | LOC4346285 | dihydroneopterin aldolase 2 | Os08g0556200 | 3.4 | 3.8 | 2.9 | 2.5 |
| | 23 | LOC4343425 | WAT1-related protein At5g64700 | Os07g0524900 | 3.3 | 2.2 | 4.5 | 4.1 |
| | 24 | LOC4344204 | momilactone A synthase-like | | 3.3 | 4.5 | 1 | 2.8 |
| | 25 | LOC4334614 | L-type lectin-domain containing receptor kinase SIT2-like | Os03g0823000 | 3.3 | 3.2 | 3.3 | 5.9 |
| | 26 | LOC4331157 | heavy metal-associated isoprenylated plant protein 3 | Os02g0818900 | 3.2 | 3.2 | 3.3 | |
| | 27 | LOC4333666 | BTB/POZ domain and ankyrin repeat-containing protein NPR3-like | Os03g0667100 | 3.2 | 2.6 | 3.9 | 2.2 |
| | 28 | LOC4330110 | uncharacterized LOC4330110 | Os02g0640300 | | | | |
| | 29 | LOC4325828 | NAC domain-containing protein 48-like | Os01g0946200 | 3.2 | 2.9 | 3.4 | 1.9 |
| | 30 | LOC4325589 | tryptophan aminotransferase-related protein 3 | Os01g0717700 | 3.1 | 4.4 | 1.8 | 1 |

| | | | | | | | | |
|---|---|---|---|---|---|---|---|---|
| | 31 | LOC4344755 | zingiberene synthase | Os08g0168000 | 3.1 | 2.4 | 3.8 | 2.2 |
| | 32 | LOC4345935 | tricin synthase 2-like | OSNPB_080498400 | 3.1 | 2.8 | 3.5 | 1.5 |
| | 33 | LOC4332755 | transcription factor WRKY45-1-like | Os03g0335200 | 3.1 | 3 | 3.2 | 1.3 |
| | 34 | LOC4333712 | uncharacterized LOC4333712 | Os03g0676400 | 3 | 1.3 | 4.6 | 1.1 |
| | 35 | LOC4347211 | probable potassium transporter 17 | OSNPB_090448200 | 3 | 2.5 | 3.5 | 3.5 |
| | 36 | LOC4339633 | mitogen-activated protein kinase 7-like | OSNPB_050566400 | 3 | 2.8 | 3.1 | 2 |
| | 37 | LOC4337975 | acyl transferase 7-like | OSNPB_050179300 | 3 | 3.1 | 2.8 | 2.2 |
| | 38 | LOC4343899 | protein TIFY 10b-like | OSNPB_070615200 | 3 | 2.8 | 3.1 | 2.3 |
| | 39 | LOC4343432 | bisdemethoxycurcumin synthase-like | Os07g0526400 | 3 | 3.9 | | 2.3 |
| | 40 | LOC4338646 | annexin D3 | Os05g0382900 | 3 | 3.2 | 2.7 | 1.9 |
| | 41 | LOC4345934 | tricin synthase 1-like | OSNPB_080498100 | 2.9 | 2.5 | 3.3 | 1.2 |
| | 42 | LOC107276160 | protein SAR DEFICIENT 1 | Os08g0360300 | 2.9 | 4.6 | 2.7 | 3 |
| | 43 | LOC4328485 | probable 4-coumarate--CoA ligase 3 | OSNPB_020177600 | 2.2 | 3.4 | 2.4 | 1.4 |
| | 44 | LOC4330037 | phenylalanine ammonia-lyase | Os02g0626600 | 4.5 | 5.2 | 3.8 | 2.5 |
| | 45 | LOC4345205 | cytosolic sulfotransferase 5 | Os08g0297800 | 2.9 | 1.8 | 3.9 | 1.4 |
| | 46 | LOC4334125 | protein STRICTOSIDINE SYNTHASE-LIKE 3 | Os03g0750700 | 2.9 | 2.6 | 3.3 | 2.3 |
| | 47 | LOC4340220 | U-box domain-containing protein 70-like | Os06g0163000 | 2.9 | 3.6 | 2.2 | 1.1 |
| | 48 | LOC4334427 | indole-3-glycerol phosphate lyase, chloroplastic | Os03g0797400 | 2.9 | 2.6 | 3.7 | 2.9 |
| | 49 | LOC4339070 | protein trichome birefringence-like 28 | Os05g0470000 | 2.9 | 3.4 | 2.5 | 1.6 |
| | 50 | LOC4338861 | phenylalanine ammonia-lyase | Os05g0427400 | 4.5 | 5.2 | 3.8 | 2.5 |

### Supplementary Table 5. Primers used in this study.

| Primers | Sequences | Purpose |
|---|---|---|
| NbBalSα-TsgA-F | ACACCTGCAACGAAACGAGCCATGGAGAGAACTCACG TTTTAGAGCTAG | For making CRISPR/Cas9 vector for *Nbbalsα* mutants |
| NbBalSα-TsgB-F | ACGTCTCGCAGGCACCTGCAACGGTGCTCGATCAACAC GGCGAGCAGGTTTTAGAGCTAG | For making CRISPR/Cas9 vector for *Nbbalsα* mutants |
| NbBalSα-TsgC-F | ACGTCTCGCAGGCACCTGCAACGCAGGGTCCCAAGCTT TCTGCACGGGTTTTAGAGCTAG | For making CRISPR/Cas9 vector for *Nbbalsα* mutants |
| NbBalSβ-QsgA-F | ACACCTGCAACGAAACGCTTACCATTTAGCTCAGAGGT TTTAGAGCTAG | For making CRISPR/Cas9 vector for *Nbbalsβ* mutants |
| NbBalSβ-QsgB-F | ACGTCTCGCAGGCACCTGCAACGGTGCgagagcggttcagtttgt caGTTTTAGAGCTAG | For making CRISPR/Cas9 vector for *Nbbalsβ* mutants |
| NbBalSβ-QsgC-F | ACGTCTCGCAGGCACCTGCAACGCGACTCTCTCAGCAC CAATGATCGGTTTTAGAGCTAG | For making CRISPR/Cas9 vector for *Nbbalsβ* mutants |
| NbBalSβ-QsgD-F | ACGTCTCGCAGGCACCTGCAACGCTGGACATAGGATAC TCATCCTCGGTTTTAGAGCTAG | For making CRISPR/Cas9 vector for *Nbbalsβ* mutants |
| NbBalR1-TsgA-F | ACACCTGCAACGAAACACACTCAAGTTGTTTCAGGAGT TTTAGAGCTAG | For making CRISPR/Cas9 vector for *Nbbalr1* mutants |
| NbBalR1-TsgB-F | CGTCTCGCAGGCACCTGCAACGGTGCCTATCCATTGGT AGGTTTTCGTTTTAGAGCTAG | For making CRISPR/Cas9 vector for *Nbbalr1* mutants |
| NbBalR1-TsgC-F | CGTCTCGCAGGCACCTGCAACGCAGGATAACTTGATTG ACTCGCCAGTTTTAGAGCTAG | For making CRISPR/Cas9 vector for *Nbbalr1* mutants |
| COMMON-FIRST-R | TCACCTGCTAGTGCACGCTCGAACACGTACCCGGCCGC GATTGGCCATAAGTAACCTTTAGAGT | Reverse primer annealed with all TsgA-F primer to generate the first sgRNA |
| TRI-sgB-R | TCACCTGCTAGTCCTGGCTCGAACACGTACCCGGCCGC GATTGGCCATAAGTAACCTTTAGAGT | Reverse primer annealed with all TsgB-F primers to generate the middle sgRNA |
| QUAD-sgB-R | TCACCTGCTAGTGTCGGCTCGAACACGTACCCGGCCGC GATTGGCCATAAGTAACCTTTAGAGT | Reverse primer annealed with NbBalSB-QsgB-F to generate NbBalSB-sgB |
| QUAD-sgC-R | TCACCTGCTAGTCCAGGCTCGAACACGTACCCGGCCGC GATTGGCCATAAGTAACCTTTAGAGT | Reverse primer annealed with NbBalSB-QsgC-F to generate NbBalSB-sgC |
| COMMON-LAST-R | TCACCTGCTAGTCACTTTGGCCATAAGTAACCTTT | Reverse primer annealed with all TsgC-F and QsgD-F to generate the last sgRNA |
| BalSα-gRT1 | CCGCGAGTTCTGCCTCGACCgttttagagctagaaat | For making CRISPR/Cas9 vector for *Osbalsα* mutants |
| BalSα-OsU6aT1 | GGTCGAGGCAGAACTCGCGGCggcagccaagccagca | For making CRISPR/Cas9 vector for *Osbalsα* mutants |
| BalSα-gRT2 | TGCACGGCGGCCTCCAAGGCgttttagagctagaaat | For making CRISPR/Cas9 vector for *Osbalsα* mutants |
| BalSα-OsU6bT2 | GCCTTGGAGGCCGCCGTGCACaacacaagcggcagc | For making CRISPR/Cas9 vector for *Osbalsα* mutants |
| BalSβ-gRT1 | CCTCCACCCTCGCCACCCAgttttagagctagaaat | For making CRISPR/Cas9 vector for *Osbalsβ* mutants |
| BalSβ-OsU6aT1 | TGGGTGGCGAGGGTGGAGGCggcagccaagccagca | For making CRISPR/Cas9 vector for *Osbalsβ* mutants |
| BalSβ-gRT2 | TCTCCACGGCTGCGACGAGGgttttagagctagaaat | For making CRISPR/Cas9 vector for *Osbalsβ* mutants |
| BalSβ-OsU6bT2 | CCTCGTCGCAGCCGTGGAGACaacacaagcggcagc | For making CRISPR/Cas9 vector for *Osbalsβ* mutants |
| BalSα-KpnI-F | GGGGTACCAATGGAGAACCATGCTAAAGTGG | For making the NbBalSα complementation vector |
| BalSα-XmaI-N-R | TCCCCCCGGGCTAAAGGATGAGAAAATTGGGA | For making the NbBalSα complementation vector |
| BalSβ-SalI-F | ACGCGTCGACGATGGAAAATCGTGGAAAGAAG | For making the NbBalSβ complementation vector |
| BalSβ-XmaI-N-R | TCCCCCCGGGTCACATATATGAGCGCATACGA | For making the NbBalSβ complementation vector |
| MpBalSα-XbaI-F | gagaacacgggggactctagaATGGTGCACGACCTGACCG | For making the MpBalSα complementation vector |
| MpBalSα-SpeI-N-R | tgggtatctaggcctactagtCTACAAAGAGGACCAGACGGGG | For making the MpBalSα complementation vector |
| MpBalSβ-XbaI-F | gagaacacgggggactctagaATGGCTGCACCACCGCCG | For making the MpBalSβ complementation vector |
| MpBalSβ-SpeI-N-R | tgggtatctaggcctactagtTCACAAAAATGAACTAAGGCGGG | For making the MpBalSβ complementation vector |
| BalSα-pLou3-F | gccatggatcacgtggaattcATGGAGAACCATGCTAAAGTGGC | For making the NbBalSα protein expression vector |
| BalSα-pLou3-R | caggtcgactctagaggatccCTAAAGGGATGAGAAAATTGGGAC | For making the NbBalSα protein expression vector |
| BalSβ-pLou3-F | gccatggatcacgtggaattcATGGAAAATCGTGGAAAGAAGG | For making the NbBalSβ protein expression vector |

| | | |
|---|---|---|
| BalSβ-pLou3-R | caggtcgactctagaggatccTCACATATATGAGCGCATACGAGG | For making the NbBalSβ protein expression vector |
| BalSα-Y257A-mF | TCAGTGGTGCGTgcTTTAATCCATGACT | For making the NbBalSα protein expression vector |
| BalSα-Y257A-mR | TCATGGATTAAAGCACGCACCACTGAAGTC | For making the NbBalSα protein expression vector |
| BalSα-E264A-Y265A-mF | CATGACTCTTCAGcAgcTGTTTCAGGTAACATT | For making the NbBalSα protein expression vector |
| BalSα-E264A-Y265A-mR | TACCTGAAACAGCTGCTGAAGAGTCATGGATT | For making the NbBalSα protein expression vector |
| BalSβ-ET24-F | taagaaggagatatacatatgGAAAATCGTGGAAAGAAGGTGTTG | For making the NbBalSβ protein expression vector in pull-down assay |
| BalSβ-ET24-R | acggagctcgaattcggatccTCACATATATGAGCGCATACGAGG | For making the NbBalSβ protein expression vector in pull-down assay |
| TDNA-LP | CGCTTTCCATCTAGCCAAAC | For confirming Arabidopsis T-DNA line |
| TDNA-RP | GAGCCATCGCTGATCAAGTAG | For confirming Arabidopsis T-DNA line |
| WiscDs_LB | TCCTCGAGTTTCTCCATAATAATGT | For confirming Arabidopsis T-DNA line |
| BalSα-eGFP-KpnI-F | tgcccgactatgccgggtaccGGGAGAACCATGCTAAAGTGGC | For making EGFP fusion construct |
| BalSα-eGFP-KpnI-R | ctgggacgtcataggggtaccCTAAAGGGATGAGAAAATTGGGAC | For making EGFP fusion construct |
| BalSβ-eGFP-KpnI-F | tgcccgactatgccgggtaccGGGAAAATCGTGGAAAGAAGGTG | For making EGFP fusion construct |
| BalSβ-eGFP-KpnI-R | ctgggacgtcataggggtaccTCACATATATGAGCGCATACGAGG | For making EGFP fusion construct |
| BalR1-eGFP-KpnI-F | tgcccgactatgccgggtaccGGGCAAAAACAACCCAAAACC | For making EGFP fusion construct |
| BalR1-eGFP-KpnI-R | ctgggacgtcataggggtaccTTAAAGTTTTGATGATTGACCAGCA | For making EGFP fusion construct |
| nYFP-BalSα- KpnI-F | gtctatatcatggccggtaccGAGAACCATGCTAAAGTGGCTACC | For making nYFP- and cYFP fusion construct |
| nYFP-BalSα-SpeI-R | tgggtatctaggcctactagtCTAAAGGGATGAGAAAATTGGGAC | For making nYFP- and cYFP fusion construct |
| cYFP-BalSβ- KpnI-F | gacgagctgtacaagggtaccGAAAATCGTGGAAAGAAGGTGTTG | For making nYFP- and cYFP fusion construct |
| cYFP-BalSβ-SpeI-R | tgggtatctaggcctactagtTCACATATATGAGCGCATACGAGG | For making nYFP- and cYFP fusion construct |
| ADH1-PDK1-SacI-F | gagagaacagaattcgagctcGTCATTCGACTTCATGATTAACACTG | For making RNAi vector |
| ADH1-PDK1-KpnI-R | ttccttaccaattggggtaccTAATTGACATCCTTCTTTATTAACCTTTC | For making RNAi vector |
| ADH1-PDK2-BamHI-F | gaaatcgataagcttggatccTAATTGACATCCTTCTTTATTAACCTTTC | For making RNAi vector |
| ADH1-PDK2-XbaI-R | cttgcatgcctgcagtctagaGTCATTCGACTTCATGATTAACACTG | For making RNAi vector |
| ADH3-PDK1-SacI-F | gagagaacagaattcgagctcGAAAGTGTCGGTGAAGGAGTGAC | For making RNAi vector |
| ADH3-PDK1-KpnI-R | ttccttaccaattggggtaccAAGAAGGCAGACTTTCTCCAGAGG | For making RNAi vector |
| ADH3-PDK2-BamHI-F | gaaatcgataagcttggatccAAGAAGGCAGACTTTCTCCAGAGG | For making RNAi vector |
| ADH3-PDK2-XbaI-R | cttgcatgcctgcagtctagaGAAAGTGTCGGTGAAGGAGTGAC | For making RNAi vector |
| ADH4-PDK1-SacI-F | gagagaacagaattcgagctcAGATATGAAAGCAGGAGATCGCG | For making RNAi vector |
| ADH4-PDK1-KpnI-R | ttccttaccaattggggtaccGTGGAAACGCCACAGCTAAGC | For making RNAi vector |
| ADH4-PDK2-BamHI-F | gaaatcgataagcttggatccGTGGAAACGCCACAGCTAAGC | For making RNAi vector |
| ADH4-PDK2-XbaI-R | cttgcatgcctgcagtctagaAGATATGAAAGCAGGAGATCGCG | For making RNAi vector |
| ADH7-PDK1-SacI-F | gagagaacagaattcgagctcGTGGAGAGTATTGGAGAAGATGTTCA | For making RNAi vector |
| ADH7-PDK1-KpnI-R | ttccttaccaattggggtaccGAGGCAAGCCCTGTTAGGTGG | For making RNAi vector |
| ADH7-PDK2-BamHI-F | gaaatcgataagcttggatccGAGGCAAGCCCTGTTAGGTGG | For making RNAi vector |
| ADH7-PDK2-XbaI-R | cttgcatgcctgcagtctagaGTGGAGAGTATTGGAGAAGATGTTCA | For making RNAi vector |
| *NbL06g04700.1*-RT-F | GCTGGGTATGTCATTCGACTT | Quantitative real-time PCR |
| *NbL06g04700.1*-RT-R | CGATGCTCTTCATACCGAGAA | Quantitative real-time PCR |
| *NbL06g17510.1*-RT-F | ATCCACTCGTCCATTTCAGC | Quantitative real-time PCR |
| *NbL06g17510.1*-RT-R | TGTCCGCAAGCGTCATATT | Quantitative real-time PCR |
| *NbL16g26700.1*-RT-F | ATGAACGACCGCCAGAGTC | Quantitative real-time PCR |
| *NbL16g26700.1*-RT-R | AAACAGCTCCAAGGCCAGT | Quantitative real-time PCR |
| *NbL10g24620.1*-RT-F | GAAGGAGCACGAATCAGAGG | Quantitative real-time PCR |
| *NbL10g24620.1*-RT-R | CCCTTATTTTCTCATGTACGGGTAT | Quantitative real-time PCR |
| *NbL17g27060.1*-RT-F | GTGGAAGGGGCACGAAC | Quantitative real-time PCR |
| *NbL17g27060.1*-RT-R | AATTTTCTCATGTAGGGGCATATC | Quantitative real-time PCR |
| *NbL18g12390.1*-RT-F | AATAAAACTGCTGGAAACCCAAT | Quantitative real-time PCR |
| *NbL18g12390.1*-RT-R | AAGCATGCAGGTTCTTTCAAC | Quantitative real-time PCR |
